# Phages overcome bacterial immunity via diverse anti-defense proteins

**DOI:** 10.1101/2023.05.01.538930

**Authors:** Erez Yirmiya, Azita Leavitt, Allen Lu, Carmel Avraham, Ilya Osterman, Jeremy Garb, Sadie P. Antine, Sarah E. Mooney, Sam J. Hobbs, Philip J. Kranzusch, Gil Amitai, Rotem Sorek

**Author notes:** These authors contributed equally to this work (Erez Yirmiya, Azita Leavitt).

## Abstract

It was recently shown that bacteria employ, apart from CRISPR-Cas and restriction systems, a considerable diversity of phage resistance systems, but it is largely unknown how phages cope with this multilayered bacterial immunity. Here, we analyzed groups of closely related *Bacillus* phages that showed differential sensitivity to bacterial defense systems, and identified multiple families of anti-defense proteins that inhibit the Gabija, Thoeris, and Hachiman systems. We show that these proteins efficiently cancel the defensive activity when co-expressed with the respective defense system or introduced into phage genomes. Homologs of these anti-defense proteins are found in hundreds of phages that infect taxonomically diverse bacterial species. We show that an anti-Gabija protein, denoted Gad1, blocks the ability of the Gabija defense complex to cleave phage-derived DNA. Our data further reveal an anti-Thoeris protein, denoted Tad2, which is a “sponge” that sequesters the immune signaling molecules produced by Thoeris TIR-domain proteins in response to phage. Our results demonstrate that phages encode an arsenal of anti-defense proteins that can disable a variety of bacterial defense mechanisms.

## Introduction

The arms race between bacteria and their viruses has fueled the evolution of defense systems that protect bacteria from phage infection^1–4^. Phages, in return, developed mechanisms that allow them to overcome bacterial defenses^5^. Multiple phages were shown to encode anti-restriction proteins, which inhibit restriction-modification (RM) systems by direct binding to the restriction enzyme^6, 7^ or by masking restriction sites^8^. Phages are also known to encode many CRISPR-Cas inhibitors, which function via a variety of mechanisms including inhibition of CRISPR RNA loading^9^, diversion of the CRISPR-Cas complex to bind non-specific DNA^10^, prevention of target DNA binding or cleavage^11^, and many additional mechanisms^12–15^. Phage proteins and non-coding RNAs that inhibit toxin-antitoxin-mediated defense have also been described^16, 17^.

Whereas early research focused on RM and later on CRISPR-Cas as the main mechanisms of defense against phage, recent studies exposed dozens of previously unknown defense systems that are widespread among bacteria and archaea^2, 18–23^. These systems mediate defense by employing a plethora of molecular mechanisms, including small-molecule signaling^20, 24–28^, production of antiviral compounds^22, 29^, and reverse transcription of non-coding RNAs^21, 30^. Recently, several phage proteins that inhibit the bacterial CBASS and Thoeris defense systems have been described^31–33^. However, it is still mostly unknown whether and how phages can overcome the wide variety of newly reported defense systems. In the current study we rely on comparative genomics of closely related phages to discover phage proteins that inhibit the Gabija, Thoeris, and Hachiman defense systems.

## Results

### Identification of phage genes that inhibit bacterial defenses

In a previous study, we demonstrated that analysis of genomically-similar phages that display differential sensitivity to bacterial immunity enabled the discovery of a phage protein, called Tad1, that inhibits the Thoeris defense system^33^. To examine whether this methodology could be used to systematically detect anti-defense proteins within phages, we isolated and analyzed several groups of closely related phages, and tested their sensitivity to multiple defense systems that protect *Bacillus* species from phage infection^18^ (fig. 1a). One group included eight newly isolated phages from the SPbeta group, which also includes the previously isolated phages SPbeta, phi3T and SPR^34–36^. These are temperate *Siphoviridae* phages with ∼130kb genomes, with 43%-96% alignable genomes when comparing phage pairs from this group (ED fig. 1a, Supplementary tables 1,2). A second group included six phages similar to phage SPO1, a lytic *Myoviridae* phage with a ∼130kb-long genome^37^. Over 85% of the genome was alignable when comparing phage pairs from this group, with high (80%-99%) sequence identity in alignable regions (fig. 1b, Supplementary tables 3,4). The third group of phages included eight previously-isolated phages from the SBSphiJ group, which were reported in a recent study^33^ (ED fig. 1b, Supplementary tables 5,6).

**Figure 1.**
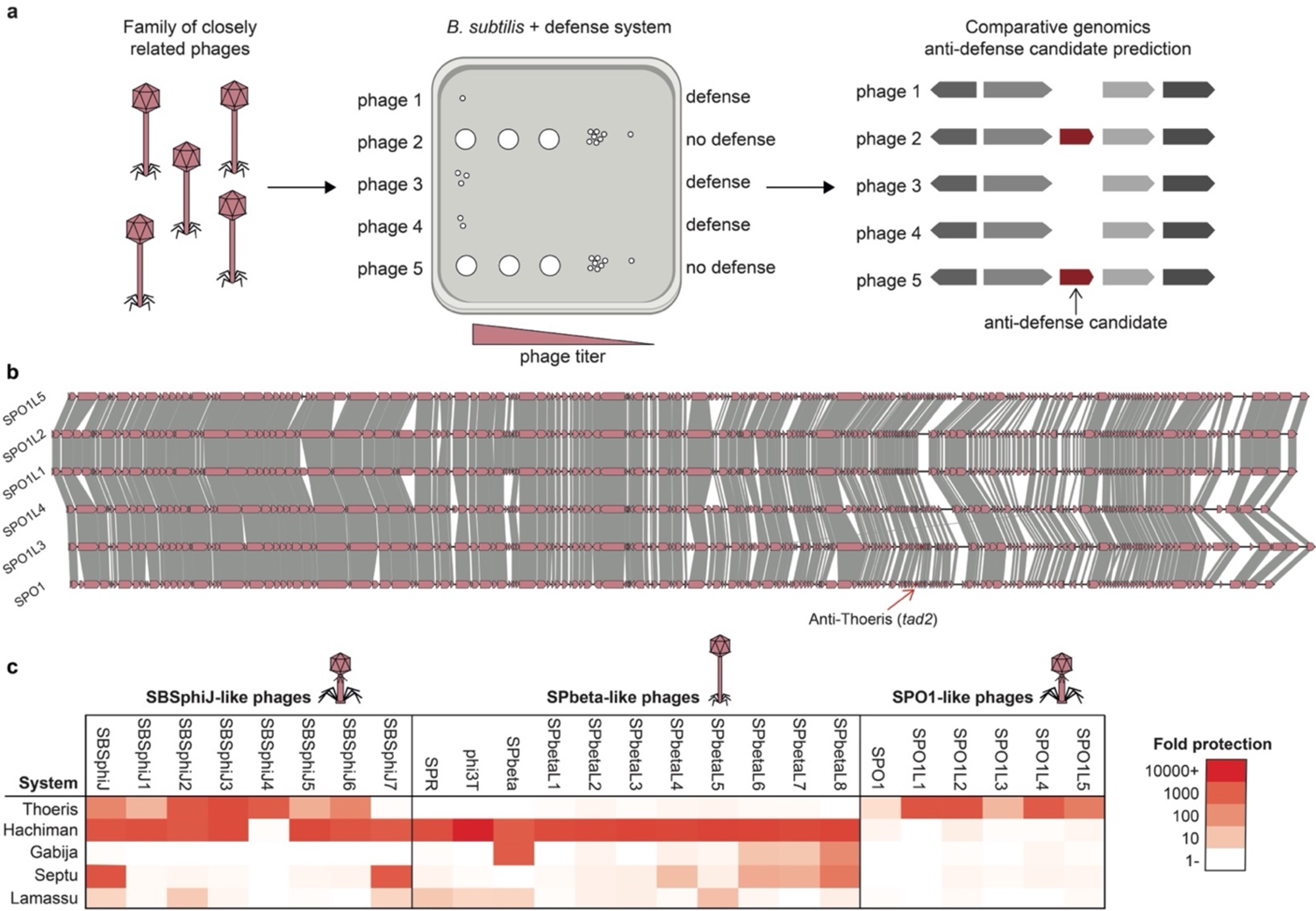
Identification of anti-defense genes based on differential sensitivity to defense systems. (a) A flowchart of the experiments and analyses used in this study to detect candidate anti-defense genes. (b) Genome comparison of six phages from the SPO1 group. Amino acid sequence similarity is marked by grey shading. Genome similarity was visualized using clinker^38^. (c) Infection profile of SBSphiJ-like, SPbeta-like and SPO1-like phages infecting five *Bacillus subtilis* strains that express each of the defense systems Thoeris, Hachiman, Gabija, Septu and Lamassu. Fold defense was measured using serial dilution plaque assays, comparing the efficiency of plating (EOP) of phages on the system-containing strain to the EOP on a control strain that lacks the systems and contains an empty vector instead. Data represent an average of three replicates. Detailed data from individual plaque assays are found in ED fig. 2.

We next used this set of 25 phages to infect strains of *Bacillus subtilis* that expressed each of the defense systems described in Doron *et al*^18^ as protecting against phages in *B. subtilis.* Five of these systems protected against at least one of the phages tested (fig. 1c). However, phages from the same group displayed remarkably different properties when infecting defense-system-containing bacteria. For example, the Gabija defense system provided strong protection against phages SPbeta and SPbetaL8 but not against other phages from the SPbeta group such as SPR and phi3T; and the Hachiman defense system provided defense against all phages of the SBSphiJ group except SBSphiJ4 (fig. 1c).

To identify phage genes that may explain the differential defense phenotype, we analyzed the gene content in groups of phages that overcame each defense system and compared it to the gene content in phages that were blocked by the system. Genes common to phages that overcame the defense system, which were not found in any of the phages that were blocked by the defense system, were considered as candidate anti-defense genes and were further examined experimentally.

### Phage genes that inhibit the Gabija defense system

Gabija is a widespread bacterial defense system found in >15% of sequenced bacterial and archaeal genomes^39^. This system comprises two genes, *gajA* and *gajB*, which encode an ATP-dependent DNA endonuclease and a UvrD-like helicase domain, respectively^18, 40^. The Gabija system was shown to provide defense against a diverse set of phages^18, 41^.

The Gabija system from *Bacillus cereus* VD045, when cloned within *B. subtilis*, provided strong protection against some phages of the SPbeta group including SPbeta and SPbetaL8, and intermediate, weaker defense against phages SPbetaL6 and SPbetaL7 (fig. 1c, ED fig. 2a). The remaining seven phages of the SPbeta group were able to completely overcome Gabija-mediated defense. We found two genes that were present in the seven Gabija-overcoming phages while missing from phages that were sensitive to Gabija defense (fig. 2a, ED fig. 1a, Supplementary table 7). One of these genes did not show a Gabija-inhibiting phenotype when co-expressed with Gabija, and we were unable to clone the second gene into Gabija-expressing cells, presumably due to toxicity. To examine the possible function of the second, non-cloned gene, we knocked out that gene from the genome of phage phi3T. Our results show that phi3T knocked out for the candidate gene was no longer able to overcome Gabija defense, suggesting that this gene inhibits Gabija (fig. 2b). We denote the anti-Gabija gene *gad1* (Gabija anti-defense 1). Engineering *gad1* from phi3T together with its native promoter into the genome of phage SPbeta, which naturally lacks this gene, rendered SPbeta resistant to Gabija, confirming that *gad1* is both necessary and sufficient for the anti-Gabija phenotype (fig. 2b).

**Figure 2.**
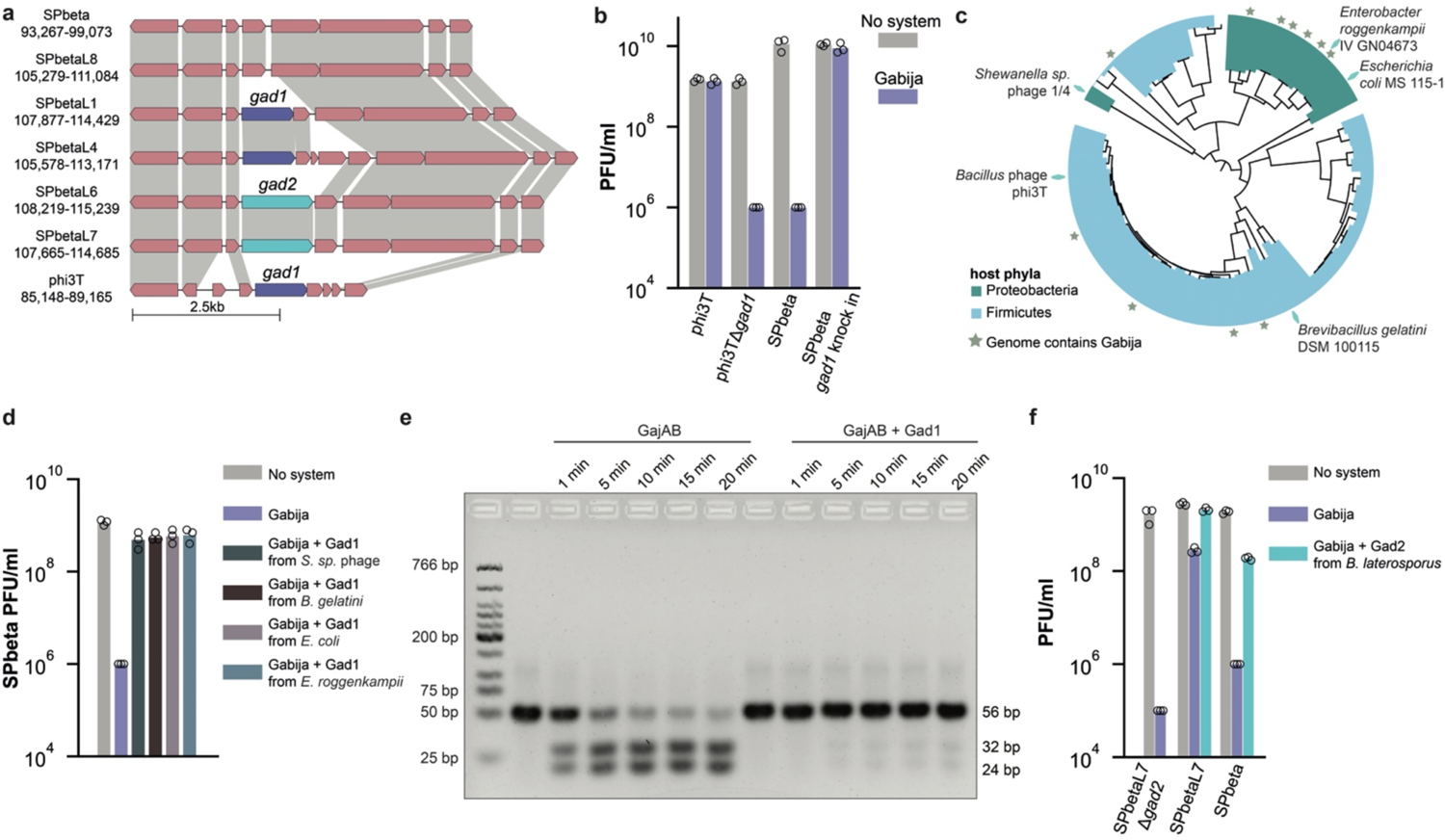
Gad1 and Gad2 proteins inhibit Gabija defense. (a) The anti-Gabija locus in phages of the SPbeta group. Amino acid sequence similarity is marked by grey shading. Genome similarity was visualized using Clinker^38^. (b) Deletion of *gad1* from phage phi3T eliminates the ability of the phage to cancel Gabija-mediated defense, while knock in of *gad1* into phage SPbeta renders the phage resistant to Gabija. Data represent plaque-forming units per ml (PFU/ml) of phages infecting control cells (“no system”) and cells expressing the Gabija defense system. Shown is the average of three replicates, with individual data points overlaid. (c) Phylogeny and distribution of Gad1 homologs. The names of bacteria in which Gad1 homologs were found in prophages and verified experimentally are indicated on the tree by cyan diamonds. (d) Results of phage infection experiments. Data represent plaque-forming units per ml (PFU/ml) of SPbeta infecting control cells (“no system”), cells expressing the Gabija system (“Gabija”), and cells co-expressing the Gabija system and a Gad1 homolog. Shown is the average of three replicates, with individual data points overlaid. (e) Gad1 blocks Gabija-mediated DNA cleavage. Incubation of purified Gabija (GajAB) complex, or Gabija co-purified with Gad1 from *Shewanella sp.* phage 1/4 (GajAB + Gad1) with a previously described DNA substrate from phage Lambda^40^. Shown is representative agarose gel from three independent experiments of proteins with 1, 5, 10, 15 or 20 min incubation with DNA. (f) Knockout of *gad2* from phage SPbetaL7 renders the phage sensitive to Gabija defense, while expression of a Gad2 homolog from a *Brevibacillus laterosporus* prophage allows SPbeta to overcome Gabija-mediated defense. Phage infection experiments were conducted as in panel (d).

Gad1 is a 295aa-long protein, which does not exhibit sequence similarity to proteins of known function. We found 94 homologs of Gad1, distributed in genomes of various phages and prophages infecting host bacteria from the phyla *Proteobacteria* and *Firmicutes* (fig. 2c, Supplementary tables 8,9). Interestingly, many of the Gad1-contaning prophages were integrated in bacterial genomes that also encoded the Gabija system, suggesting that Gad1 enabled these phages to overcome Gabija-mediated defense of their hosts (fig. 2c). Five Gad1 homologs from phages infecting *Shewanella sp.*, *Enterobacter roggenkampii*, *Escherichia coli*, *Brevibacillus gelatini* and *Bacillus xiamenensis* were selected for further experimental examination (fig. 2c, ED fig. 3a). Unlike the Gad1 protein from phage phi3T, four of the Gad1 homologs were not toxic when expressed in Gabija-containing cells (all except the homolog cloned from the *B. xiamenensis* prophage). Each of these four homologs efficiently inhibited Gabija defense when co-expressed in Gabija-containing cells (fig. 2d, ED fig. 3b).

**Figure 3.**
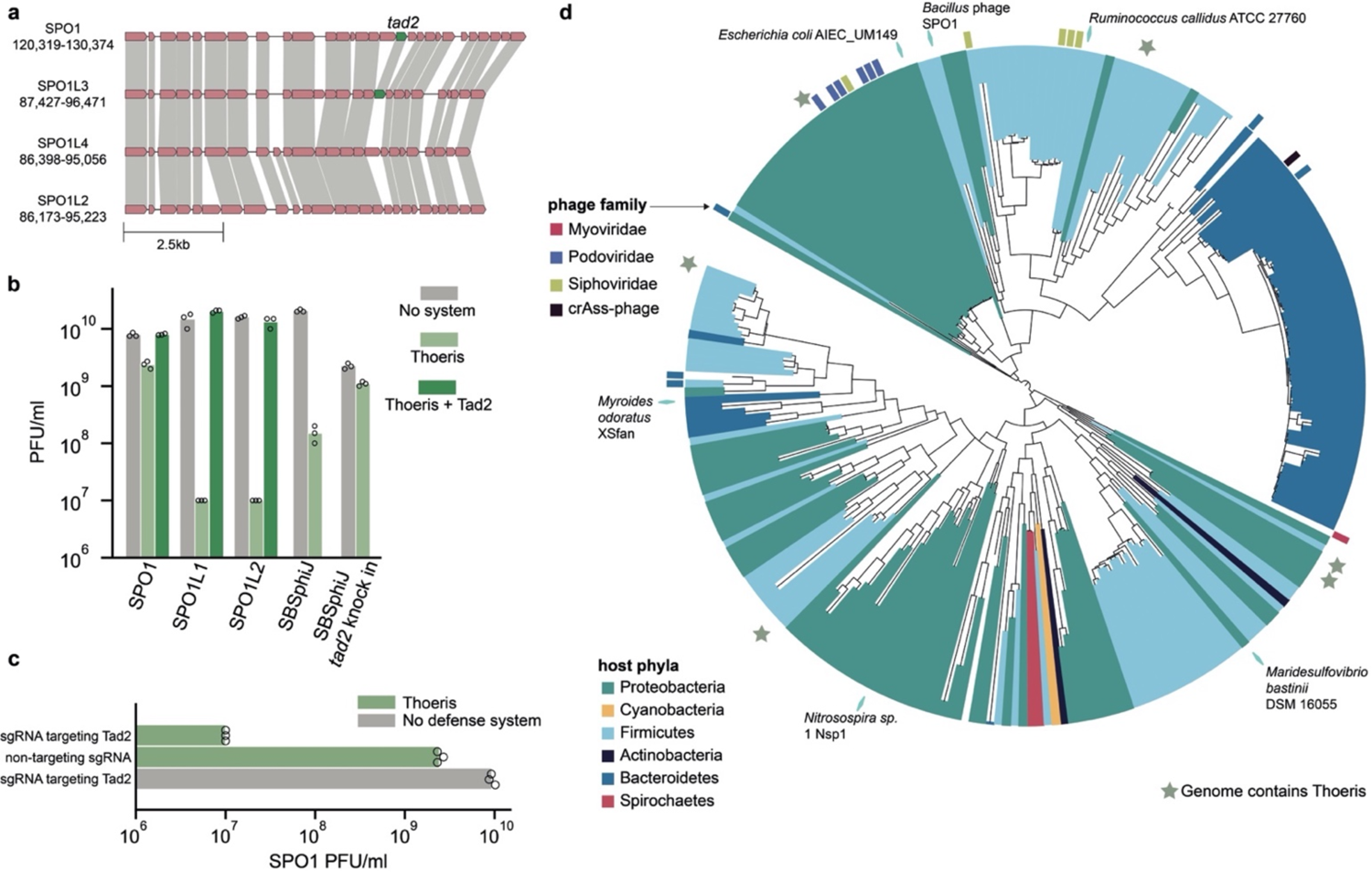
Tad2 proteins inhibit Thoeris defense. (a) Genomic locus of the anti-Thoeris gene *tad2* in phages SPO1 and SPO1L3, and the respective locus in phages SPO1L4 and SPO1L2. Amino acid sequence similarity is marked by grey shading. Genome similarity was visualized using Clinker^38^. (b) Anti-Thoeris activity of Tad2. Data represent plaque-forming units per milliliter (PFU/ml) of phages infecting control cells (no system), cells expressing the Thoeris system (Thoeris) and cells co-expressing the Thoeris system and the *tad2* gene from SPO1 (Thoeris + Tad2). Data for phage SBSphiJ as well as for SBSphiJ with a *tad2* knock in are also presented. Shown is the average of three replicates, with individual data points overlaid. (c) Tad2 knockdown cancels anti-Thoeris activity. Data represent PFU/ml of SPO1 that infects cells expressing Thoeris and a dCas9 system targeting Tad2, as well as control cells. (d) Phylogenetic analysis of Tad2 homologs in phage and prophage genomes. The names of bacteria in which Tad2 homologs were found in prophages and tested experimentally are indicated on the tree by cyan diamonds.

It was recently shown that Gabija identifies and cleaves DNA having a specific nearly palindromic sequence motif derived from phage Lambda^40^. By purifying GajA and GajB and reconstituting the Gabija complex *in vitro*, we were able to confirm that Gabija cleaves DNA that contains the specific sequence motif (fig. 2e). Gabija purified in the presence of Gad1 was unable to cleave DNA (fig. 2e). In a companion paper, we show that Gad1 binds the Gabija complex as an octamer and inhibits its ability to bind and cleave DNA (Antine et al 2023, submitted manuscript).

We next examined phages SPbetaL6 and SPbetaL7, which lack *gad1* but were still partially resistant to Gabija (fig. 1c). Intriguingly, these phages encoded, at the same locus where *gad1* was encoded in other phages, another gene of unknown function. Knocking-out this gene from phage SPbetaL7 rendered this phage completely sensitive to Gabija (fig. 2f). The gene from SPbetaL7 was toxic when expressed in bacteria, but co-expression of a non-toxic homolog from a prophage of *Brevibacillus laterosporus* with Gabija completely inhibited Gabija defense, further verifying that it is an anti-Gabija gene which we denote here *gad2* (fig. 2a,f). Gad2 is a 400aa-long protein that shows no sequence similarity to Gad1 or to other known proteins. Homology searches identified 170 homologs of Gad2 which almost always reside in genomes of phages and prophages infecting diverse host bacteria (ED fig. 3c, Supplementary tables 10,11). Intriguingly, structural analysis using Alphafold2^42^ predicted that Gad2 is an enzyme with a nucleotidyltransferase protein domain, suggesting that it inhibits Gabija via a mechanism of action different than Gad1 (ED fig. 3d). These results suggest that SPbeta-like phages encode anti-Gabija genes in a dedicated locus in their genomes, where multiple different Gabija-inhibiting genes can reside.

### Phage genes that inhibit the Thoeris defense system

The Thoeris defense system is present in approximately 4% of sequenced bacterial and archaeal genomes^18^. This system encodes ThsB, a protein with a Toll/interleukin-1 receptor (TIR) domain that serves as a sensor for phage infection, and ThsA, an NAD^+^-cleaving protein^27^. Upon phage recognition, the Thoeris ThsB protein generates 1′′–3′ gcADPR, a signaling molecule that activates ThsA, resulting in the depletion of NAD^+^ and the inhibition of phage replication^33^.

Our recent discovery of the anti-Thoeris gene *tad1*, present in phage SBSphiJ7 and absent from other phages in the SBSphiJ group, explains the observed insensitivity of SBSphiJ7 to Thoeris^33^ (fig. 1c, ED fig. 2c). We hypothesized that phages SPO1 and SPO1L3 may also encode homologs of *tad1*, as these phages partially escaped Thoeris-mediated defense (fig. 1c, ED fig. 2b), but we were unable to identify *tad1* homologs in these phages. Instead, we identified a single gene present in SPO1 and SPO1L3 but absent from all other Thoeris-sensitive SPO1-like phages (fig. 3a, Supplementary table 7). We expressed this gene, designated here *tad2,* within *B. subtilis* cells that also express the Thoeris system from *B. cereus* MSX-D12. Tad2 robustly inhibited the activity of Thoeris, allowing Thoeris-sensitive phages to infect Thoeris-expressing cells (fig. 3b). Moreover, engineering *tad2* into SBSphiJ, a phage that is normally blocked by Thoeris, resulted in a phage that overcomes Thoeris-mediated defense (fig. 3b). Silencing the expression of Tad2 in SPO1 using dCas9 ^43^ further confirmed that Tad2 is responsible for the Thoeris-inhibiting phenotype of SPO1 (fig. 3c).

Tad2 is a short protein (89 amino acids) containing a DUF2829 protein domain. Sequence homology searches identified over 650 direct homologs in the integrated microbial genomes (IMG)^44^ and metagenomic gut virus (MGV)^45^ databases (Supplementary tables 12,13). Phylogenetic analysis revealed that Tad2 is encoded by phages belonging to several phage morphology groups, including *Myoviridae*, *Podoviridae* and *Siphoviridae*, as well as by prophages integrated within over 100 bacterial species from 6 different phyla (fig. 3d). We selected 5 Tad2 homologs representing the phylogenetic diversity of the family and cloned each one separately into *B. subtilis* cells expressing the Thoeris system (fig. 3d, ED fig. 4a). Four of these Tad2 homologs were able to inhibit Thoeris, including homologs derived from phages infecting distant organisms such as *Ruminococcus callidus* and *Maridesulfovibrio bastinii* (ED fig. 4b). These results demonstrate that Tad2 represents a large family of proteins used by phages to inhibit the Thoeris defense system.

**Figure 4.**
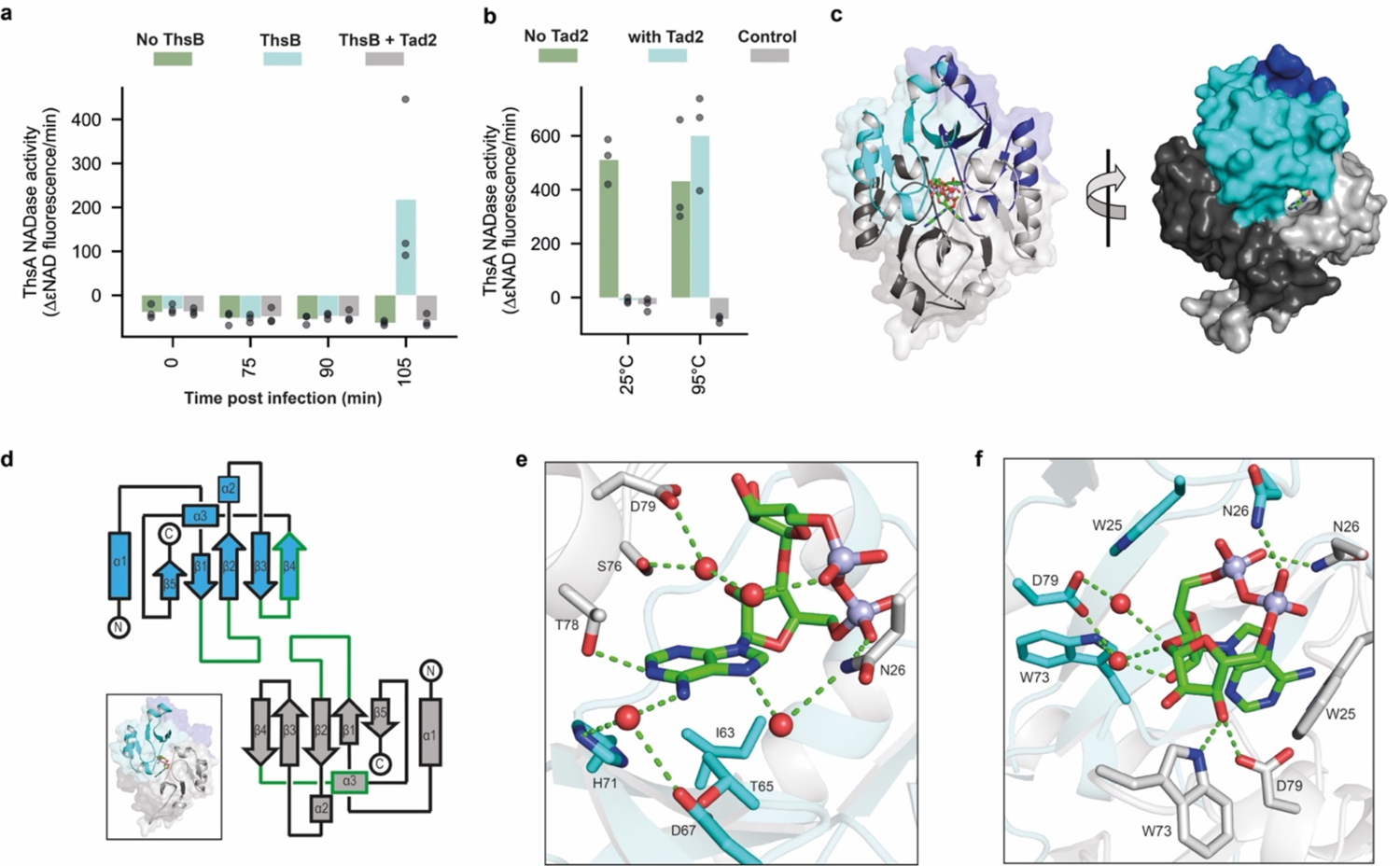
Tad2 cancels Thoeris-mediated defense by sequestering 1ʹʹ–3ʹ gcADPR. (a) Cells expressing ThsB, both ThsB and Tad2 or control cells that do not express ThsB were infected with phage SBSphiJ at a multiplicity of infection (MOI) of 10. NADase activity of purified ThsA incubated with filtered lysates was measured using a nicotinamide 1,N6-ethenoadenine dinucleotide (εNAD) cleavage fluorescence assay. Bars represent the mean of three experiments, with individual data points overlaid. (b) Tad2 releases bound 1ʹʹ–3ʹ gcADPR when denatured. Shown is the NADase activity of purified ThsA incubated with 600nM 1’’-3’ gcADPR preincubated with 2.4 μM of purified Tad2 *in vitro* for 10 min, followed by an additional incubation of 10 min at either 25 °C or 95 °C. Control is buffer without 1ʹʹ–3ʹ gcADPR. (c) Overview of the crystal structure of Tad2 in complex with 1ʹʹ–3ʹ gcADPR in cartoon (front) or surface (side) representation. Tad2 exists as a homotetramer formed by two dimer units (colored cyan/dark blue and grey/dark grey). Non-dimeric monomer pairs form two recessed ligand binding pockets that enclose 1ʹʹ–3ʹ gcADPR. (d) Topology map of two Tad2 monomers which come together to form the ligand binding pocket. Components that form the binding pocket are outlined in green. Each dimer subunit donates one monomer, as shown by the cartoon representation. (e,f) Detailed views centered around the adenine base (e) or ribose and phosphates (f) of residues that either directly interact with 1ʹʹ–3ʹ gcADPR or coordinate key waters that reside within the binding pocket. Residues contributed by each of the two monomers that form the binding pocket are represented in cyan (Tad2a) or grey (Tad2b).

During phage infection, ThsB generates the 1′′–3′ gcADPR signaling molecule to activate ThsA^33^. As expected, purified ThsA incubated with filtered cell lysates derived from SBSphiJ-infected, ThsB-expressing cells became a strong NADase, indicating that ThsB produced 1′′–3′ gcADPR in response to SBSphiJ infection as previously shown^33^ (fig. 4a). However, filtered lysates from similarly infected cells that co-expressed both ThsB and Tad2 failed to activate ThsA *in vitro*, suggesting that Tad2 functions upstream of ThsA (fig. 4a).

Recent studies have shown that phages can use dedicated proteins to degrade^32, 46^ or sequester^31, 33^ bacterial immune signaling molecules. Incubation of Tad2 with 1′′–3′ gcADPR *in vitro* did not yield observable degradation products, suggesting that Tad2 is not an enzyme that cleaves 1′′–3′ gcADPR (ED fig. 5). To test whether Tad2 might act as a sponge that binds and sequesters the immune signal, we incubated purified Tad2 with 1′′–3′ gcADPR, and then boiled the reaction in 95 °C to denature Tad2. The boiled reaction readily activated ThsA, suggesting that Tad2 functions by binding and sequestering the Thoeris-derived signaling molecule, and that denaturation of Tad2 released the intact molecule back to the medium (fig. 4b). In support of this observation, Tad2 that was pre-incubated with 1′′–3′ gcADPR showed a substantial mobility shift during size-exclusion chromatography (ED fig. 6a). In addition, Tad2 exhibited increased absorption ratio of UV_260nm_/UV_280nm_ following incubation with 1′′–3′ gcADPR, further confirming that Tad2 binds this molecule as a ligand (ED fig. 6a). We found that Tad2 binds 1′′–3′ gcADPR with K_D_ = 23.3 nM (ED fig. 6b). Notably, Tad2 could not bind the canonical cADPR, demonstrating high specificity of Tad2 to the ThsB-derived signaling molecule (ED fig. 6c).

**Figure 5.**
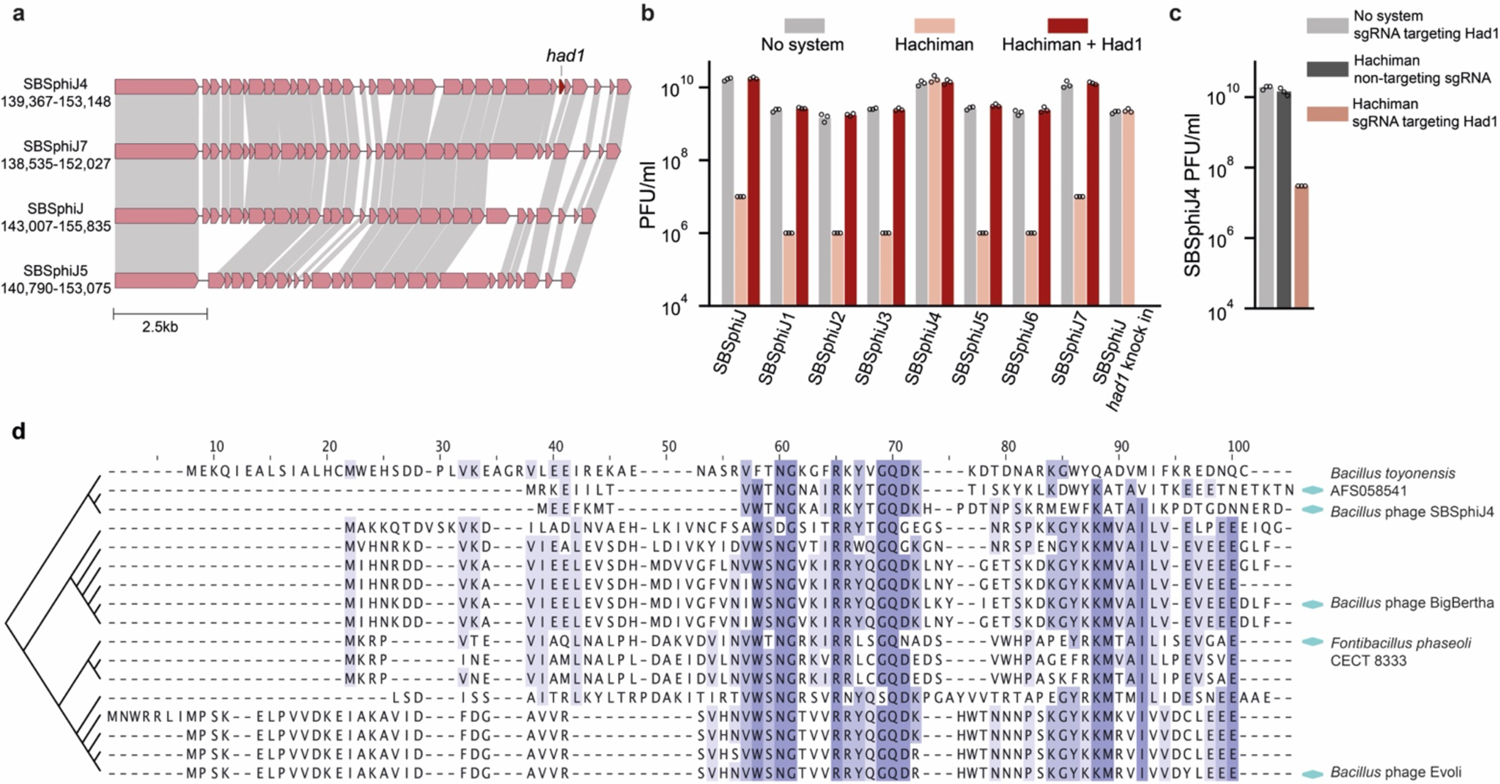
Had1 proteins inhibit Hachiman defense. (a) Genomic locus of the anti-Hachiman gene *had1* in phages SBSphiJ4, as well as the relevant locus in phage SBSphiJ7, SBSphiJ and SBSphiJ5. Amino acid sequence similarity is marked by grey shading. Genome similarity was visualized using clinker^38^. (b) Differential defense of Hachiman against phages from the SBSphiJ group, and anti-Hachiman activity of Had1. Data represent plaque-forming units per milliliter (PFU/ml) of phages infecting control cells (no system), cells expressing the Hachiman system (Hachiman) and cells co-expressing the Hachiman system and the *had1* gene from SBSphiJ4 (Hachiman + Had1). Data for phage SBSphiJ with a *had1* knock-in are also presented. Shown is the average of three replicates, with individual data points overlaid. (c) Had1 knockdown cancels anti-Hachiman activity. Results of phage SBSphiJ4 infection experiments. Data represent plaque-forming units per ml (PFU/ml) of SBSphiJ4 infecting cells expressing Hachiman and a dCas9 with an sgRNA targeting Had1, as well as control cells. Shown is the average of three replicates, with individual data points overlaid. (d) Cladogram and multiple sequence alignment of Had1 homologs. Conserved residues are colored purple. Homologs tested experimentally are indicated with cyan diamonds.

To define the mechanism of 1′′–3′ gcADPR sequestration and Thoeris inhibition, we determined crystal structures of Tad2 from phage SPO1 in the apo and 1′′–3′ gcADPR ligand-bound states (ED Table 1). The apo structure of Tad2 reveals a homotetrameric assembly consistent with oligomerization observed during size-exclusion chromatography analysis (ED fig. 7). Each Tad2 monomer contains an N-terminal alpha helix (α1) followed by an antiparallel beta sheet (β1–β5), as well as extended loops between β2–β3 and β4–β5 which contain short alpha helices α2 and α3 (fig. 4c,d). Tad2 protomers interact along two interfaces to form the tetrameric complex (fig. 4c and ED fig. 7). First, two Tad2 protomers pack together at helix α2 and sheet β2 to form V-shaped homodimeric units. Two V-shaped units interlock perpendicularly along helix α1 to complete tetramerization (fig. 4c). The resulting assembly creates two identical ligand binding pockets formed at the interface of two adjacent non-dimeric protomers, surrounded by loop β1–β2, loop β3–β4, and strand β4 of one protomer, and loop β1–β2, loop β4–α3, and helix α3 of the other (fig. 4c–f).

A 1.75 Å structure of Tad2 in complex with 1′′–3′ gcADPR explains the molecular basis of signal recognition. In the Tad2–1′′–3′ gcADPR complex, ligand binding is mediated by extensive van der Waals interactions from W25_a,b_ and W73_a,b_, as well as polar interactions to phosphates and free hydroxyls from N26_a,b_ and D79_a,b_ (fig. 4e,f). Unlike Tad1, Tad2 forms very few contacts to the adenine base of 1′′–3′ gcADPR, with one hydrogen bond from T78_b_ and nonpolar interactions from I63_a_ and T65_a_ (fig 4e,f, ED fig. 8). Additionally, we observed several key water molecules that contribute to 1′′–3′ gcADPR binding and are coordinated by N26_b_, D67_a_, H71_a_, W73_b_, S76_b_, and D79_a,b_ (fig. 4e,f). Taken together, our findings show that Tad2 inhibits Thoeris defense by binding and sequestering 1′′–3′ gcADPR, preventing the activation of the Thoeris immune effector and mitigating Thoeris-mediated defense.

### Phage genes that inhibit the Hachiman defense system

Hachiman is a defense system whose mechanism of action remains unsolved. It encodes a protein with a predicted helicase domain, as well as an additional protein with no known functional domains. Hachiman is widely distributed in genomes of bacterial and archaeal species, and was shown to provide strong protection against a broad range of phages^18^.

We cloned three short genes that were unique to phage SBSphiJ4, a phage that overcame Hachiman-mediated defense (fig. 1c, Supplementary table 7), into a *B. subtilis* strain that expresses the Hachiman system from *B. cereus* B4087 ^18^. One of these genes completely abolished Hachiman-mediated defense, and we therefore named it *had1* (Hachiman anti-defense 1) (fig. 5a,b). Silencing of Had1 expression in SBSphiJ4 resulted in a phage that could infect control strains, but was blocked by Hachiman defense (fig. 5c). In addition, SBSphiJ that was engineered to include Had1 with its native promoter was able to overcome Hachiman defense, demonstrating that Had1 is responsible for the anti-Hachiman phenotype (fig. 5b).

Had1 is a short protein sized 53 aa, which does not show sequence homology to any protein of known function. We found 23 homologs of Had1 in *Bacillus* phages as well as prophages integrated within *Bacillus* and *Paenibacillus* genomes (Supplementary table 14), and selected five homologs that span the protein sequence diversity of Had1 for further experimental examination (fig. 5d). Four of these proteins efficiently inhibited the activity of Hachiman, and we could not clone the fifth into Hachiman-expressing cells, possibly due to toxicity (fig. 5d, ED fig. 9). These results confirm that Had1 is a Hachiman-inhibiting family of phage proteins. As the mechanism of Hachiman defense is unknown, understanding how Had1 inhibits Hachiman in future studies may assist in solving the mechanism of Hachiman defense.

## Discussion

In this study we identified multiple families of phage anti-defense proteins using comparative analysis of closely-related phage genomes. Our findings demonstrate that phages have evolved diverse strategies to counter the complex, multilayered bacterial defense arsenal. Intriguingly, our data suggest that SPbeta-like phages store anti-Gabija genes in a dedicated anti-Gabija locus in their genomes. This is reminiscent of anti-CRISPR loci identified in phages of *Pseudomonas* and other species, where different sets of anti-CRISPR genes are present in each individual phage^47^. Our discovery of a specific anti-Gabija locus in SPbeta-like phages may point to a general rule for the organization of anti-defense genes in phage genomes, a genomic phenomenon that can assist in future searches for anti-defense genes.

The most widely distributed family of anti-defense proteins discovered here is that of Tad2, a protein that contains a domain of unknown function DUF2829. A protein with DUF2829 was previously speculated to inhibit type II CRISPR-Cas systems and specifically Cas9 ^48^, although binding to Cas9 was not demonstrated and the mechanism through which this protein antagonizes type II CRISPR-Cas was not determined^48^. Our findings that multiple DUF2829-containing proteins antagonize the Thoeris defense system by specifically binding and sequestering 1’’-3’ gcADPR suggest that the DUF2829 family of proteins represent anti-Thoeris proteins rather than anti-CRISPR proteins.

Our results on Tad2 join recent discoveries of additional anti-defense proteins that function by sequestering immune signaling molecules^31, 33^. These include Tad1, a completely different family of Thoeris-inhibiting proteins, which, similar to Tad2, also bind and sequester gcADPR molecules; and the phage-encoded Acb2 protein that inhibits CBASS defense by binding the cyclic oligonucleotide immune signaling molecules produced by CBASS^31, 33^. It therefore appears that the production of protein “sponges” that sequester immune signaling molecules is a highly efficient strategy that evolved within phages multiple times in parallel to allow successful evasion of immune systems that utilize immune signaling molecules. The efficiency of this strategy may be perceived as counter-intuitive, because it necessitates a 1:1 ratio between the number of phage sponge proteins and the immune signaling molecule (or even 2:1 in the case of Tad2). However, as Thoeris and CBASS become active and produce the immune signaling molecule relatively late in the infection cycle of the phage^20, 27^, phages have sufficient time to express a substantial amount of their sponge proteins at the early stages of infection. Thus, when the immune signaling molecule is produced by the defense system, there will already be enough copies of the sponge protein in the infected cell to efficiently block immune signaling.

We were not able to identify inhibitors of the Septu or Lamassu defense systems among the three groups of phages that we studied, although phages from these groups displayed differential sensitivity to these defense systems. It is possible that escape from these two systems is mediated by mutations in existing genes rather than the presence of a dedicated anti-defense gene^41^. It is also possible that different phages in the same group utilize more than one protein to overcome defense, which would sometime hamper the analysis pipeline used in this study.

With the emergence of multiple antibiotic-resistant bacteria^49^, phage therapy, in which phages are used as an alternative to antibiotics, is being considered as a suitable therapeutic avenue for defeating bacterial pathogens^50–52^. One of the major obstacles for successful phage therapy is the recently discovered ability of bacteria to actively defend themselves by encoding a large variety of defense systems. Indeed, it was shown that the set of defense systems carried by a given bacterial strain is a strong determinant for the susceptibility of that strain to phages^53, 54^. Engineering phages to carry a set of anti-defense proteins can enable them to overcome bacterial defenses, resulting in phages with increased host ranges that will be more suitable for phage therapy. Thus, the anti-defense proteins we discovered here, as well as additional such proteins that were discovered and will be discovered in the future, could be used as tools for more efficient phage therapy applications.

## Data Availability

Data that support the findings of this study are available within the article and its Supplementary Tables. IMG/MGV accessions, protein sequences and nucleotide sequences appear in Supplementary Tables 8–14. Coordinates and structure factors of cbTad1–1′′–3′-gcADPR, SPO1 Tad2 apo, SPO1 Tad2–1′′–3′-gcADPR, and SPO1 Tad2–1′′–2′-gcADPR have been deposited in the PDB under the accession codes 8SMD, 8SME, 8SMF, and 8SMG, respectively. Source data are available for all the main figures and for extended data figures 2,3,4,6,7 and 9.

## Supplementary Materials

## Extended Data Figures

**Extended Data Figure 1.**
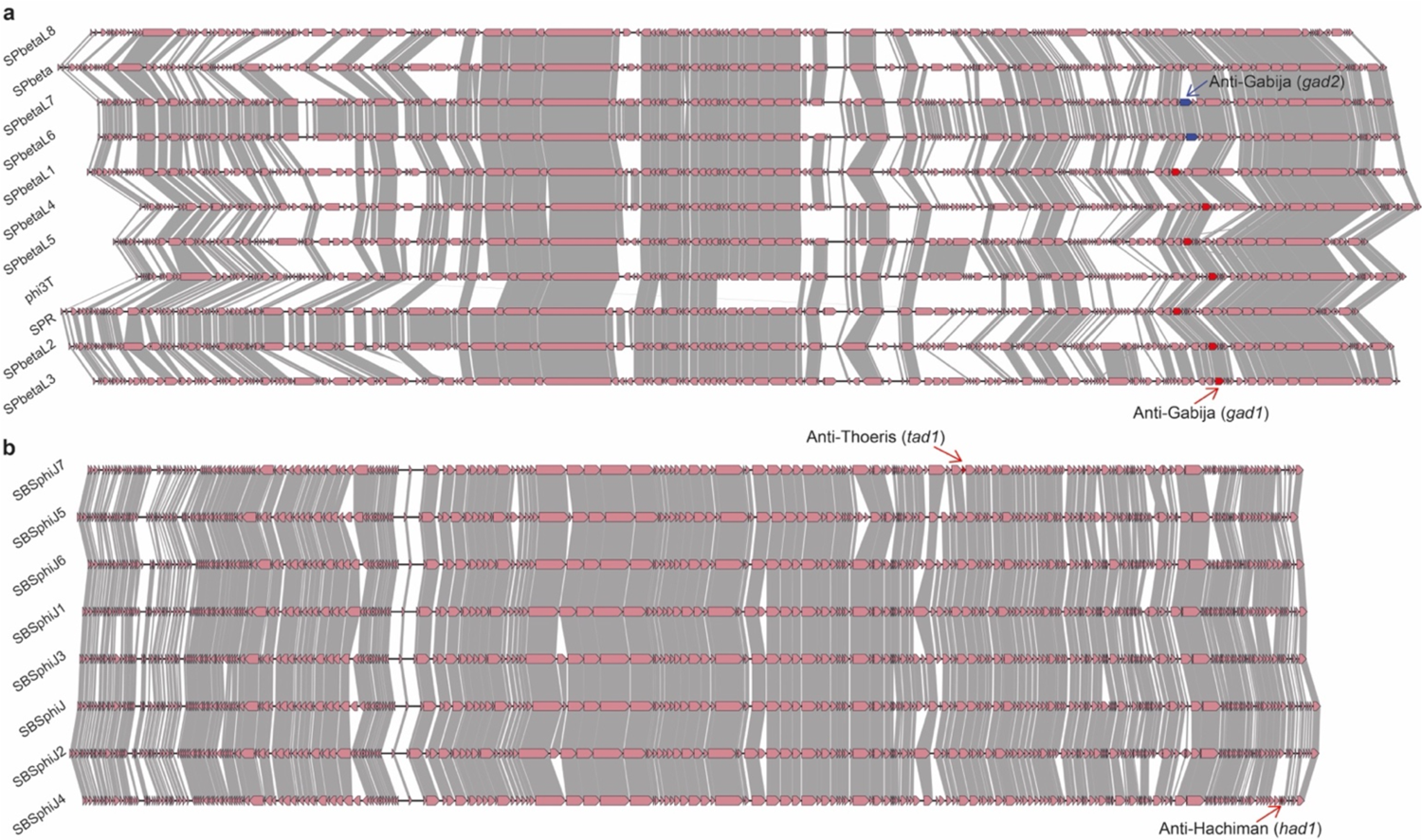
Genome comparisons of phages from the SPbeta and SBSphiJ groups. Genome comparison of (a) eleven phages from the SPbeta group and (b) eight phages from the SBSphiJ group. Amino acid sequence similarity is marked by grey shading. Genome similarity was visualized using clinker^38^.

**Extended Data Figure 2.**
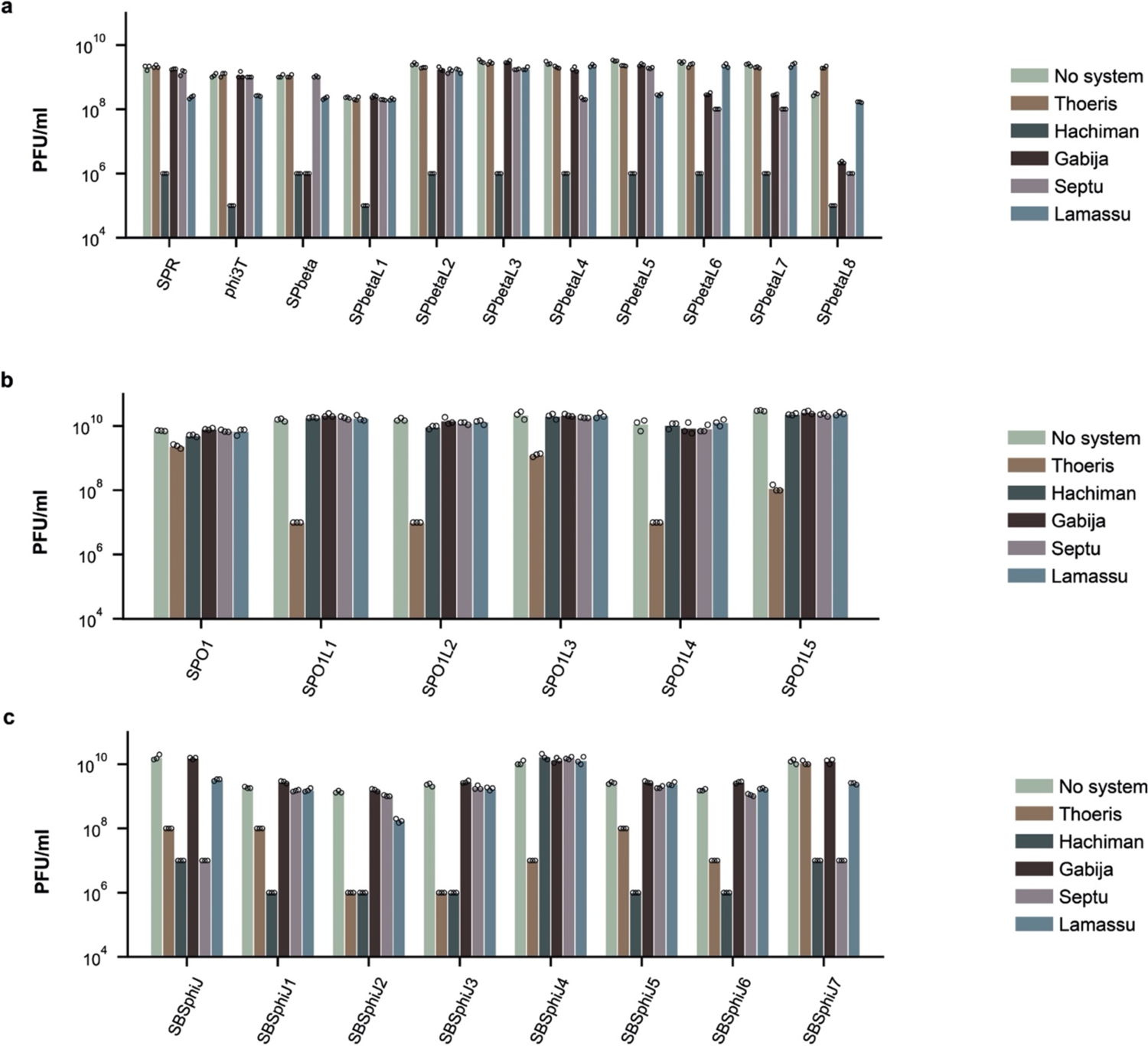
Phages from the same family are differentially sensitive to bacterial defense systems. Results of phage infection experiments with (a) eleven phages of the SPbeta group, (b) six phages of the SPO1 group, and (c) eight phages of the SBSphiJ group. Data represent plaque-forming units per ml (PFU/ml) of phages infecting control cells (“no system”), and cells expressing the respective defense systems. Shown is the average of three replicates, with individual data points overlaid. The Thoeris and Hachiman data presented here are the same as those presented in Figures 3b and 5b, respectively.

**Extended Data Figure 3.**
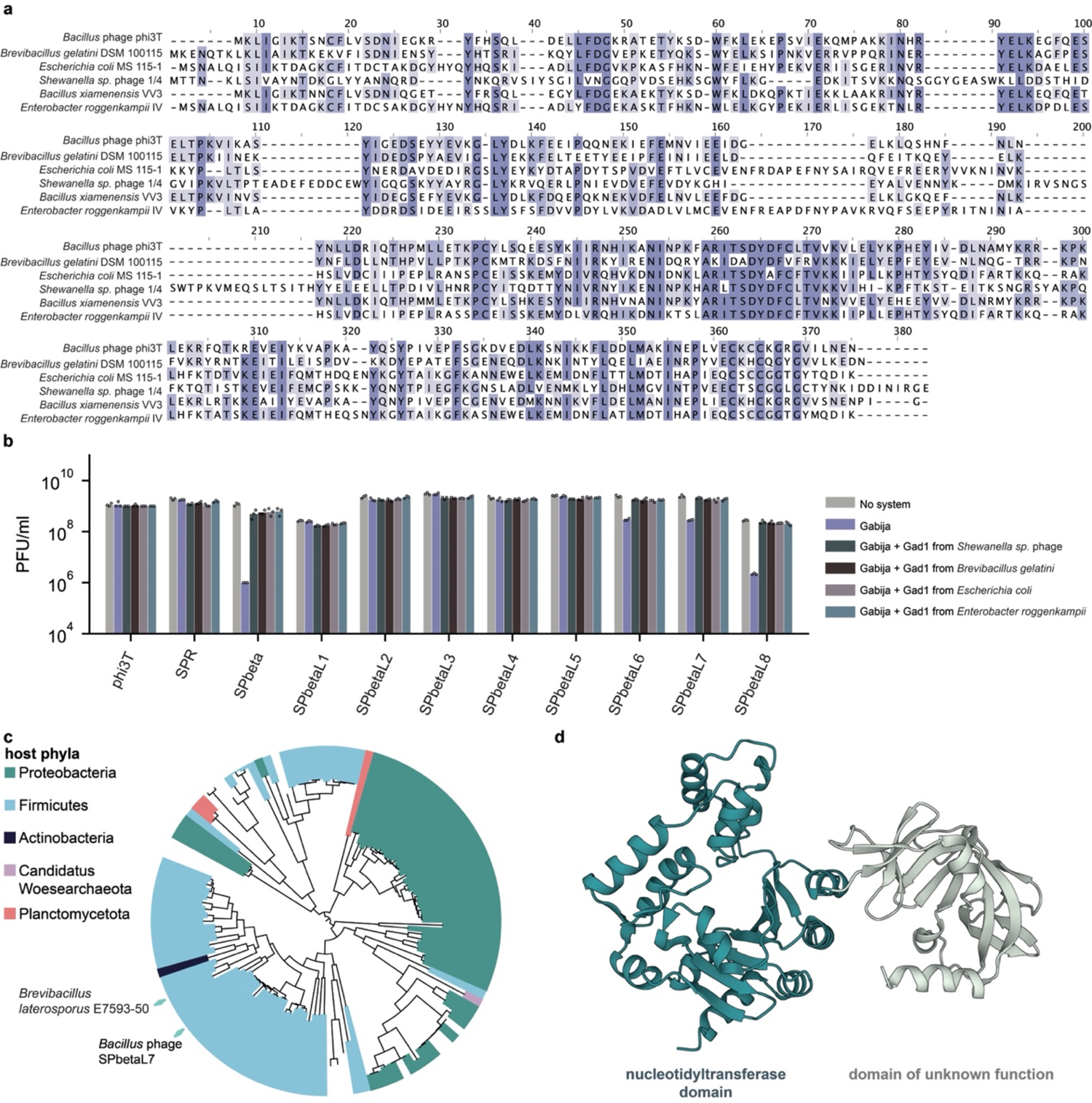
Gad1 and Gad2 proteins inhibit Gabija mediated defense. (a) Multiple sequence alignment of the original Gad1 from phage phi3T and five Gad1 homologs that were chosen for experimental verification. Conserved residues are in purple. (b) Results of phage infection experiments with eleven phages of the SPbeta group. Data represent plaque-forming units per ml (PFU/ml) of phages infecting control cells (“no system”), cells expressing the Gabija system (“Gabija”), and cells co-expressing the Gabija system and a Gad1 homolog. Shown is the average of three replicates, with individual data points overlaid. The SPbeta data presented here are the same as those presented in Figure 2d. (c) Phylogeny and distribution of Gad2 homologs. Homologs that were tested experimentally are indicated on the tree by cyan diamonds. (d) An Alphafold2 model for the structure of Gad2 from phage SPbetaL7.

**Extended Data Figure 4.**
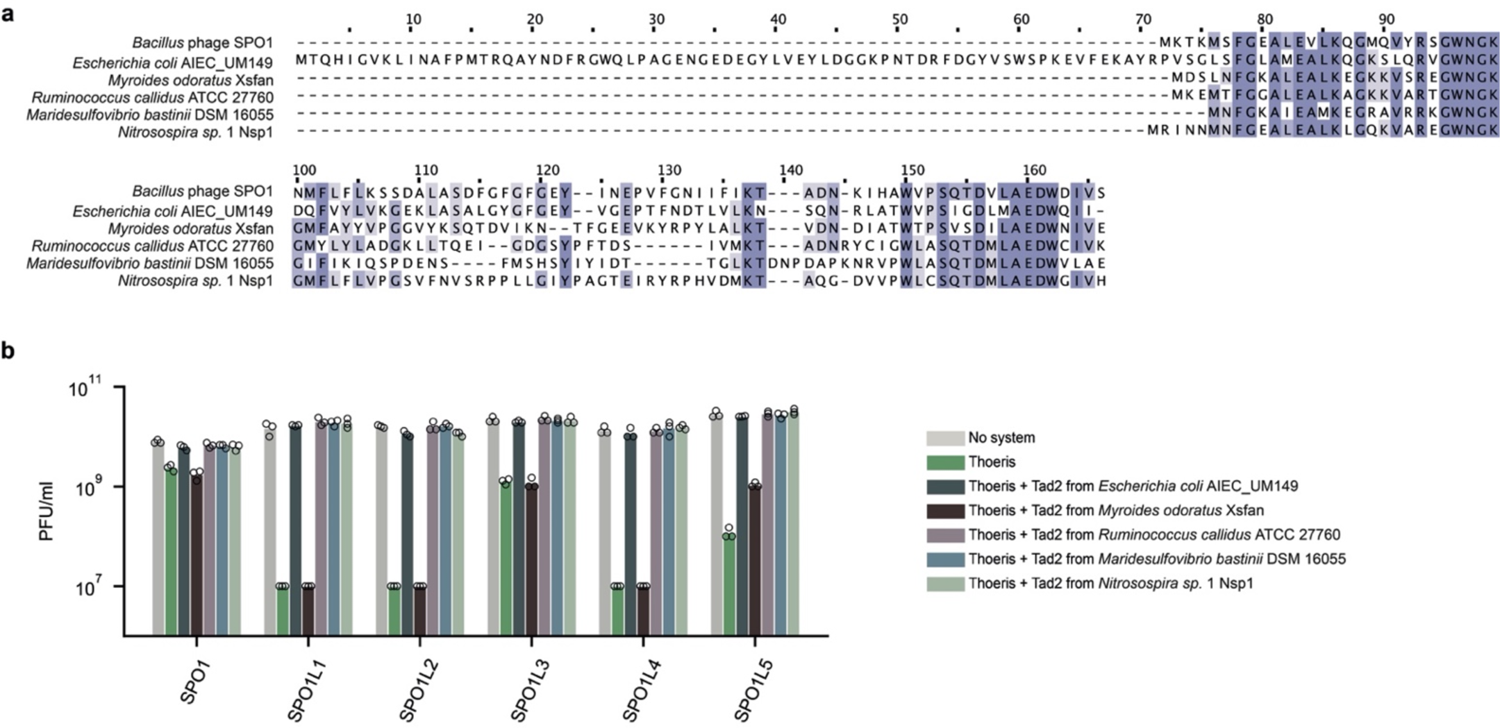
Tad2 proteins inhibit Thoeris mediated defense. (a) Multiple sequence alignment of the original Tad2 from phage SPO1, and 5 Tad2 homologs that were chosen for experimental verification. Conserved residues are in purple. (b) Results of phage infection experiments with six phages of the SPO1 group. Data represent plaque-forming units per ml (PFU/ml) of phages infecting control cells (“no system”), cells expressing the Thoeris system (“Thoeris”), and cells co-expressing the Thoeris system and a Tad2 homolog. Shown is the average of three replicates, with individual data points overlaid.

**Extended Data Figure 5.**
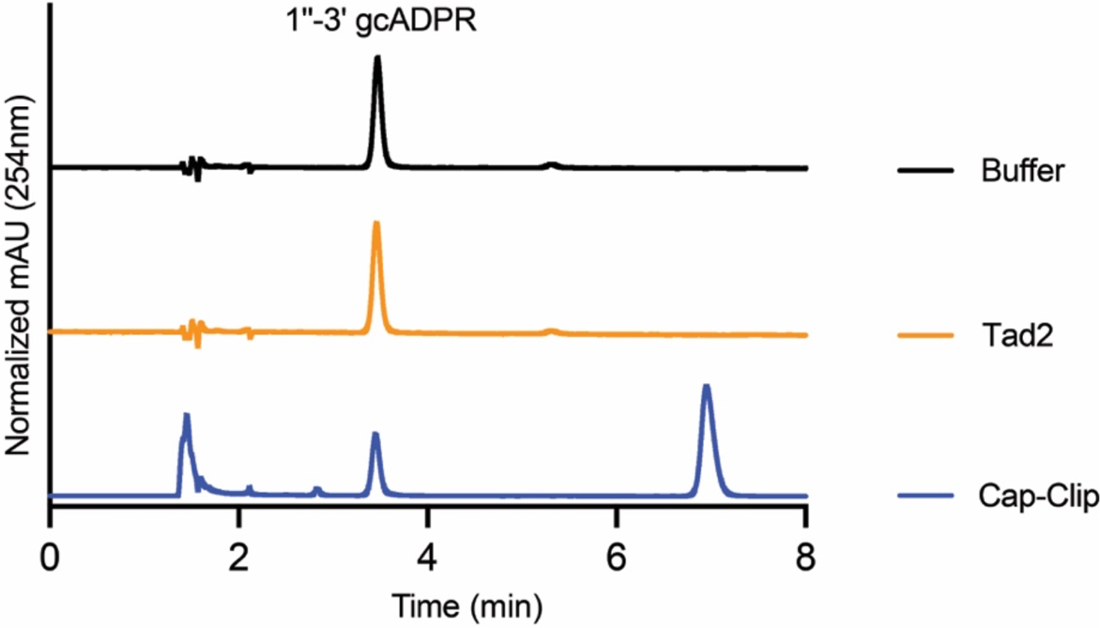
Incubation of Tad2 with 1ʹʹ–3ʹ gcADPR *in vitro* does not yield observable degradation products. Representative HPLC traces of 1ʹʹ–3ʹ gcADPR incubated with buffer, Tad2, or with the enzyme Cap-Clip known to cleave diphosphate linkages as a positive control.

**Extended Data Figure 6.**
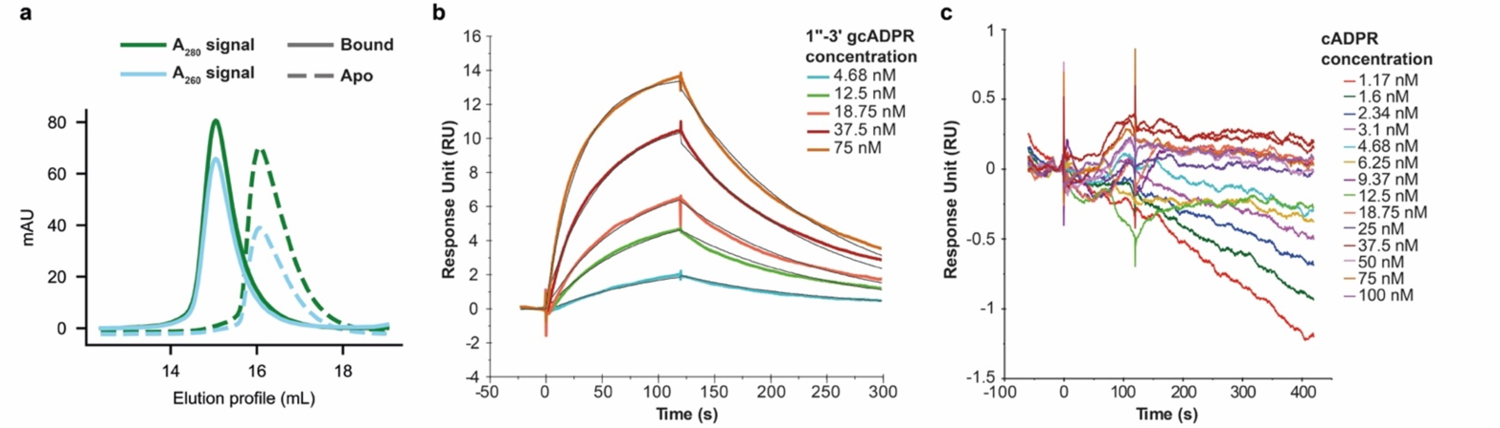
Tad2 binds 1ʹʹ–3ʹ gcADPR. (a) size-exclusion chromatography of 1ʹʹ–3ʹ gcADPR-bound or apo state Tad2. 1ʹʹ–3ʹ gcADPR-bound Tad2 shows a substantial shift compared to Tad2 in the apo state. (b) Surface plasmon resonance binding sensorgrams for Tad2 at five concentrations of 1ʹʹ–3ʹ gcADPR. The black lines are the global fits using the instrument’s built-in function. (c) Surface plasmon resonance binding sensorgrams for Tad2 at multiple concentrations of cADPR.

**Extended Data Figure 7.**
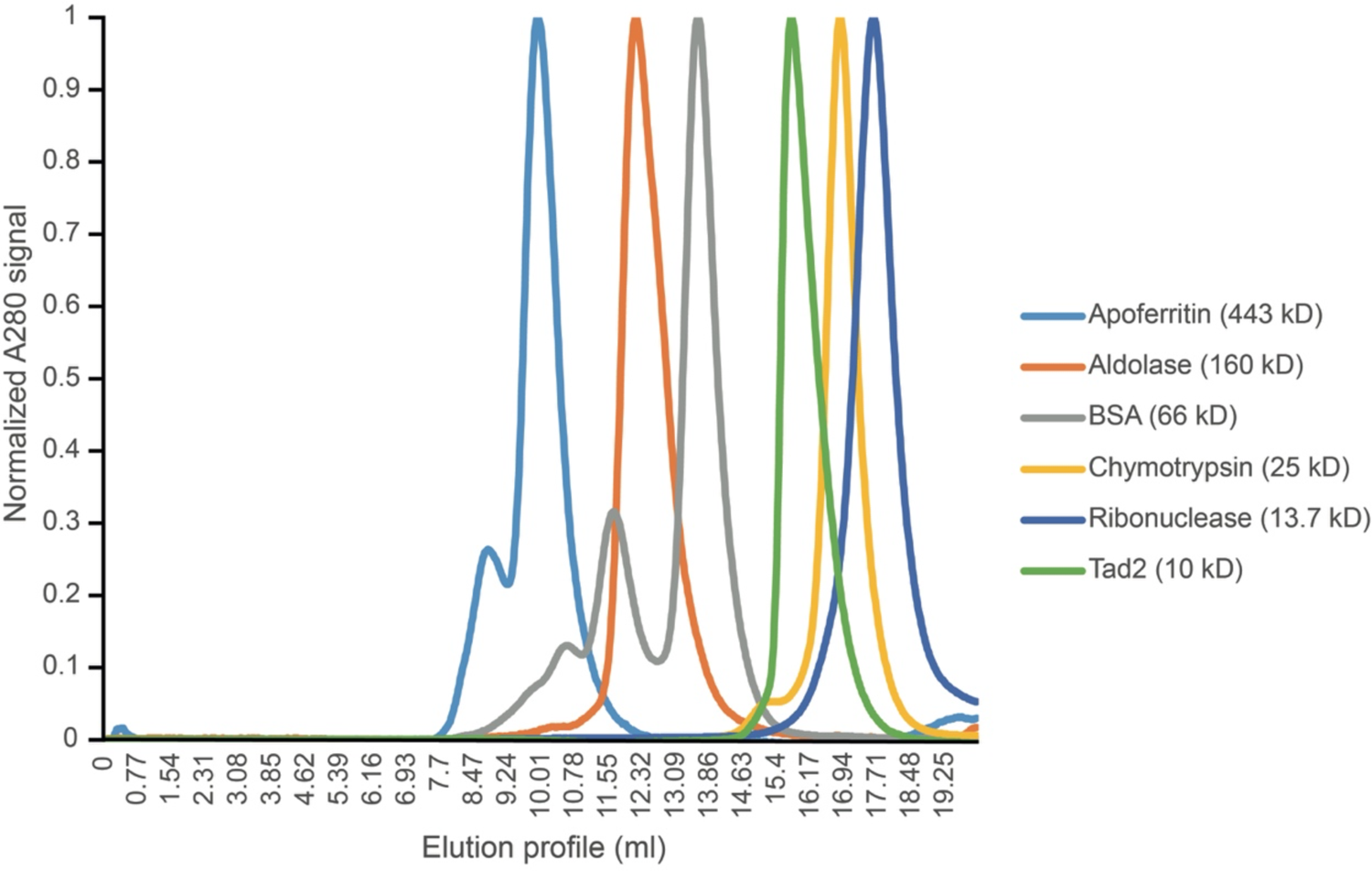
Size-exclusion chromatography of Tad2 and various standards.

**Extended Data Figure 8.**
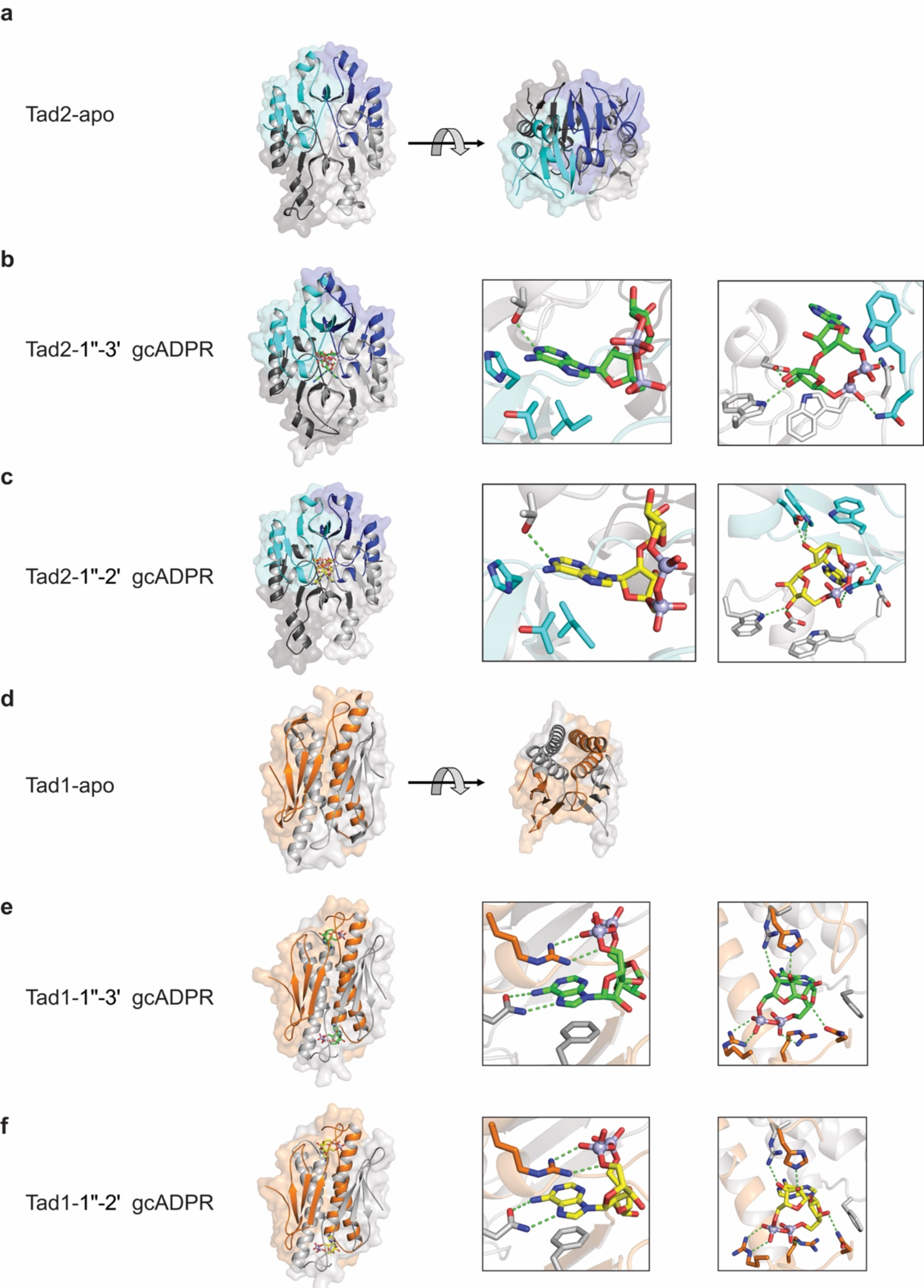
Comparison of Tad2 and Tad1 in the apo and ligand-bound states. (a) Overview of the crystal structure of SPO1 Tad2 in the apo state in front and top view. (b,c) Overview and detailed binding pocket views of adenine interactions (left) and ribose/phosphate interactions (right) of the crystal structures of SPO1 Tad2 in complex with 1ʹʹ–3ʹ gcADPR (**b**) or 1ʹʹ–2ʹ gcADPR (**c**). (d) Overview of the crystal structure (PDB: 7UAV) of cbTad1 in the apo state in front view and top view. (e,f) Overview and detailed binding pocket views of adenine interactions (left) and ribose/phosphate interactions (right) of the crystal structures of cbTad1 in complex with 1ʹʹ–3ʹ gcADPR (**e**) or 1ʹʹ–2ʹ gcADPR (**f**, PDB: 7UAW).

**Extended Data Figure 9.**
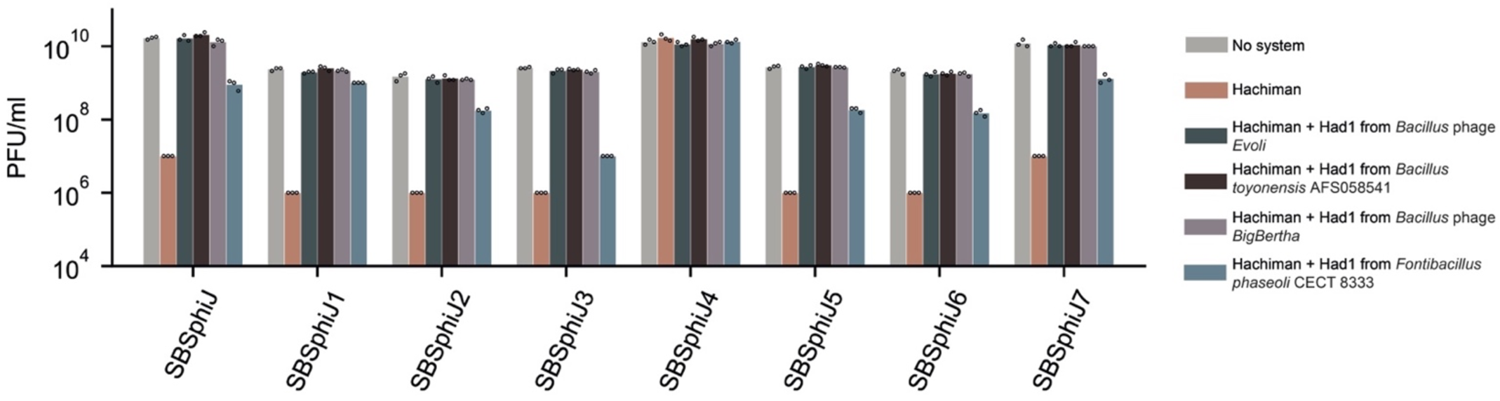
Had1 proteins inhibit Hachiman-mediated defense. Results of phage infection experiments with eight phages of the SBSphiJ group. Data represent plaque-forming units per ml (PFU/ml) of phages infecting control cells (“no system”), cells expressing the Hachiman system (“Hachiman”), and cells co-expressing the Hachiman system and a Had1 homolog. Shown is the average of three replicates, with individual data points overlaid.

## Extended Data Tables

**Extended Data Table 1:** Summary of crystallography data collection, phasing, and refinement statistics.

|  | cbTAD1–<br>1''–3' gcADPR<br>(8SMD) | SPO1 TAD2–<br>apo<br>(8SME) | SPO1 TAD2–<br>1''–3' gcADPR<br>(8SMF) | SPO1 TAD2–<br>1''–2' gcADPR<br>(8SMG) |
| --- | --- | --- | --- | --- |
| **Data collection** |  |  |  |  |
| Space group | P 2<sub>1</sub> 3 | P 6<sub>3</sub> 2 2 | C 1 2 1 | P 6<sub>3</sub> 2 2 |
| Cell dimensions |  |  |  |  |
| *a*, *b*, *c* (Å) | 95.97, 95.97,<br>95.97 | 108.40, 108.40,<br>74.26 | 94.39, 82.47,<br>90.76 | 108.06, 108.06,<br>72.36 |
| $\alpha$, $\beta$, $\gamma$ (°) | 90.00, 90.00,<br>90.00 | 90.00, 90.00,<br>120.00 | 90.00, 107.88,<br>90.00 | 90.00, 90.00,<br>120.00 |
| Resolution (Å) | 42.92–2.10<br>(2.16–2.10) | 46.94–2.36<br>(2.45–2.36) | 46.06–1.75<br>(1.78–1.75) | 39.29–2.10<br>(2.16–2.10) |
| *R*<sub>pim</sub> | 2.5 (69.8) | 1.9 (81.5) | 5.7 (63.0) | 2.2 (71.9) |
| *I* / $\sigma(I)$ | 12.2 (1.0) | 19.6 (1.5) | 6.5 (1.1) | 17.3 (1.5) |
| Completeness (%) | 99.7 (96.3) | 100.0 (100.0) | 98.8 (97.1) | 100.0 (99.8) |
| Redundancy | 13.5 (13.3) | 27.0 (26.7) | 5.2 (4.7) | 27.5 (26.8) |
| **Refinement** |  |  |  |  |
| Resolution (Å) | 42.92–2.10 | 46.94–2.36 | 46.06–1.75 | 39.29–2.10 |
| No. reflections |  |  |  |  |
| Total | 236478 | 298974 | 344731 | 414196 |
| Unique | 17522 | 11073 | 65939 | 15065 |
| Free | 876 | 1103 | 1994 | 1503 |
| *R*<sub>work</sub> / *R*<sub>free</sub> | 23.61 / 20.12 | 29.87 / 26.29 | 24.14 / 20.77 | 24.99 / 22.39 |
| No. atoms |  |  |  |  |
| Protein | 1984 (2 copies) | 1281 | 5054 (8 copies) | 1284 (2 copies) |
| Ligand / ion | 70 | – | 142 | 35 |
| Water | 60 | 2 | 599 | 27 |
| *B*-factors |  |  |  |  |
| Protein | 61.50 | 86.46 | 24.43 | 72.56 |
| Ligand / ion | 55.31 | – | 16.62 | 58.93 |
| Water | 55.46 | 59.19 | 34.43 | 59.70 |
| R.m.s. deviations |  |  |  |  |
| Bond lengths<br>(Å) | 0.001 | 0.002 | 0.004 | 0.002 |
| Bond angles (°) | 0.433 | 0.434 | 0.795 | 0.463 |

\*All datasets were collected from individual crystals. \*Values in parentheses are for the highest resolution shell.

## Supplementary Tables

**Supplementary table 1.** Percent alignable sequence between pairs of SPbeta-like phages.

**Supplementary table 2.** Percent sequence identity in alignable regions between pairs of SPbeta-like phages.

**Supplementary table 3.** Percent alignable sequence between pairs of SPO1-like phages.

**Supplementary table 4.** Percent sequence identity in alignable regions between pairs of SPO1-like phages.

**Supplementary table 5.** Percent alignable sequence between pairs of SBSphiJ-like phages.

**Supplementary table 6.** Percent sequence identity in alignable regions between pairs of SBSphiJ-like phages.

**Supplementary table 7.** Candidate anti-defense phage genes.

**Supplementary table 8.** Gad1 homologs in the IMG database.

**Supplementary table 9.** Gad1 homologs in the metagenomic gut viruses (MGV) database.

**Supplementary table 10.** Gad2 homologs in the IMG database.

**Supplementary table 11.** Gad2 homologs in the metagenomic gut viruses (MGV) database.

**Supplementary table 12.** Tad2 homologs in the IMG database.

**Supplementary table 13.** Tad2 homologs in the metagenomic gut viruses (MGV) database.

**Supplementary table 14.** Had1 homologs in the IMG database.

**Supplementary table 15.** Primers used for knockout, knockdown and knockin experiments.

## Methods

### Phage strains, isolation, cultivation and sequencing

*B. subtilis* phages phi3T (BGSCID 1L1, GenBank accession KY030782.1), SPbeta (BGSCID 1L5, GenBank accession AF020713.1), SPR (BGSCID 1L56, GenBank accession OM236515.1) and SPO1 (BGSCID 1P4, GenBank accession NC_011421.1) were obtained from the Bacillus Genetic Stock Center (BGSC). Phages from the SBSphiJ group were isolated by us in previous studies^18, 33^. Other phages used in this study were isolated by us from soil samples on *B. subtilis* BEST7003 culture as described in Doron *et al.*^18^. For this, soil samples were added to *B. subtilis* BEST7003 culture and incubated overnight to enrich for *B. subtilis* phages. The enriched sample was centrifuged and filtered through 0.45 µm filters, and the filtered supernatant was used to perform double layer plaque assays as described in Kropinski *et al*.^55^. Single plaques that appeared after overnight incubation were picked, re-isolated 3 times, and amplified as described below.

Phages were propagated by picking a single phage plaque into a liquid culture of *B. subtilis* BEST7003 grown at 37°C to OD_600_ of 0.3 in magnesium manganese broth (MMB) (LB + 0.1 mM MnCl_2_ + 5 mM MgCl_2_) until culture collapse. The culture was then centrifuged for 10 min at 3200 *g* and the supernatant was filtered through a 0.45 µm filter to get rid of remaining bacteria and bacterial debris.

High titer phage lysates (>10^7^ pfu ml^−1^) were used for DNA extraction. 500 µl of the phage lysate was treated with DNase-I (Merck cat #11284932001) added to a final concentration of 20 mg ml^−1^and incubated at 37°C for 1 hour to remove bacterial DNA. DNA was extracted using the QIAGEN DNeasy blood and tissue kit (cat #69504) starting from the Proteinase-K treatment step.

Phages from the SBSphiJ and SPbeta groups were sequenced using a modified Nextera protocol as previously described^56^. Following Illumina sequencing, adapter sequences were removed from the reads using Cutadapt version 2.8 ^57^ with the option -q 5. The trimmed reads from each phage genome were assembled into scaffolds using SPAdes genome assembler version 3.14.0 ^58^, using the –careful flag.

The genomes of phages from the SPO1 group were sequenced via a long-read PacBio method, due to the high amount of modified bases in these phages. For library construction of phages from the SPO1 group, 1 µg of genomic DNA samples were fragmented using g-tubes (Covaris). Sheared DNA was purified with AMPure PB beads and was used to construct a SMRTbell library according to the PacBio library construction guidelines^59^. Samples were barcoded using Barcoded Overhang Adapters and pooled to one final library. Quantity and quality of the SMRTbell library were determined using the Qubit HS DNA kit and Agilent TapeStation Genomic DNA. No size selection was performed. The PacBio sequencing primer was then annealed to the SMRTbell library followed by binding of the polymerase to the primer–library complex. The library was loaded onto one SMRT cell in the PacBio Sequel system and sequenced in a Continuous Long Read (CLR) mode at a 10-hour movie time. All phage genomes sequenced and assembled in this study were analyzed with Prodigal version 2.6.3^60^ (default parameters) to predict ORFs.

### Plaque assays

Phage titer was determined using the small drop plaque assay method^61^. 400 µl of overnight culture of bacteria were mixed with 0.5% agar and 30 ml MMB and poured into a 10 cm square plate followed by incubation for 1 hour at room temperature. In cases of bacteria expressing anti-defense candidates and in the experiment with phage SBSphiJ with a Tad2 knock-in, 1 mM IPTG was added to the medium. In cases of bacteria expressing dCas9-gRNA constructs, 0.002% xylose was added to the medium.10-fold serial dilutions in MMB were performed for each of the tested phages and 10 µl drops were put on the bacterial layer. After the drops had dried up, the plates were inverted and incubated at room temperature overnight. Plaque forming units (PFUs) were determined by counting the derived plaques after overnight incubation and lysate titer was determined by calculating PFUs per ml. When no individual plaques could be identified, a faint lysis zone across the drop area was considered to be 10 plaques. Efficiency of plating (EOP) was measured by comparing plaque assay results on control bacteria and bacteria containing the defense system and/or a candidate anti-defense gene.

### Prediction of candidate anti-defense genes

Predicted protein sequences from all phage genomes in each phage family were clustered into groups of homologs using the cluster module in MMSeqs2 release 12-113e3 ^62^, with the parameters -e 10, -c 0.8, -s 8, --min-seq-id 0.3 and the flag --single-step-clustering. For each defense system, anti-defense candidates were defined as clusters that have a representation in all the phages that overcome the defense system and are absent from all the phages that are blocked by the defense system. One member was chosen from each cluster as a candidate anti-defense gene for further experimental testing. In the case of the Hachiman defense system, only predicted genes with no known function were tested experimentally. *Gad2* was predicted based on its localization in the sane locus as *gad1* in phages SPbetaL6 and SPbetaL7.

### Cloning of candidate anti-defense genes

The DNA of each anti-defense candidate was amplified from the source phage genome using KAPA HiFi HotStart ReadyMix (Roche cat # KK2601). Homologs of verified anti-defense genes were synthesized by Genscript Corp. The anti-defense candidates were cloned into the pSG-*thrC*-Phspank vector^33^ and transformed into DH5α competent cells. The cloned vector was subsequently transformed into *B. subtilis* BEST7003 cells containing the respective defense systems as applicable integrated into the *amyE* locus^18^, resulting in cultures expressing both defense system and their corresponding anti-defense gene candidates. As a negative control, a transformant with an identical plasmid containing GFP instead of the anti-defense gene, was used. Transformation to *B. subtilis* was performed using MC medium as previously described^18^. Whole-genome sequencing was then applied to all transformed *B. subtilis* strains, and Breseq analysis^63^ was used to verify the integrity of the inserts and lack of mutations.

### Gad1 knockout in phi3T lysogenic strain

The upstream and downstream homologous arms of Gad1 were amplified from the phi3T phage genome using the PCR primer pairs Gad1_AF and Gad1_AR, and Gad1_BF and Gad1_BR, respectively (Supplementary table 15). The spectinomycin resistance gene cassette was amplified from the vector pJmp3 (addgene plasmid #79875) using the PCR primers Spec_F and Spec_R (Supplementary table 15). The backbone was amplified using the primers Vector_F and Vector_R (Supplementary table 15).

These three parts were assembled together with the pJmp3 backbone using the NEBuilder HiFi DNA Assembly cloning kit (NEB cat # E5520S) and transformed to DH5α competent cells. The cloned vector was then transformed into the phi3T lysogenic strain (BGSCID 1L1) and was plated on LB agar plates supplemented with 100 μg ml^−1^spectinomycin and incubated overnight at 30 °C. The modified phi3T prophage was induced using Mitomycin C (Sigma, M0503). Whole-genome sequencing was performed to verify the sequence of the modified phage.

### Gad2 knockout in SPbetaL7 using Cas13a

Cas13a was amplified from the pBA559 plasmid^64^ with the primers cas13a_fwd and cas13a_rev. The xylose promoter and homology arms for integration into the thrC site were amplified from plasmid pJG_thrC_dCAS9_gRNA^33^ with primers dcas9xylProm_fwd and dcas9xylProm_rev. The gRNA complimentary to the beginning of Gad2 was amplified from the pBA559 plasmid with primers gRNAcas13_fwd and gRNAcas13L7159_rev. Plasmid assembly was conducted using NEBuilder HiFi DNA Assembly cloning kit (NEB, cat # E5520S) and transformed to DH5α-competent cells. The cloned vector was subsequently transformed into the thrC site of *B. subtilis* BEST7003.

For the selection of Gad2 knockout SPbetaL7 phages, overnight culture of the Cas13a-gRNA containing bacteria was mixed was diluted 1:100 with MMB agar 0.5% with 0.2% xylose, and grown for 1 hour at RT. Then, 10^8^ PFU of phage SPbetaL7 were spread on the plate. On the next day, several individual plaques were collected and propagated with 1 ml of the Cas13a-gRNA containing bacteria. Gad2 knockout was verified using PCR primers L7159Fchk and L7159Rchk. Sequencing of the product demonstrated deletion of bases 109,505-110,753 from the SPbetaL7 genome, spanning the entire Gad2 gene and as well as the Gad2 promoter. The selected knockout phage was purified three times on *B. subtilis* BEST7003.

### Gabija / Gabija + Gad1 complex assembly and *in vitro* nuclease activity

*Bc*GajAB and *Bc*GajAB + *Shewanella sp*. phage 1/4 Gad1 complexes were purified as described (Antine et al. 2023 submitted manuscript). A 56-bp dsDNA with a sequence specific motif derived from phage Lambda (5′-TTTTTTTTTTTTTTTTTAATAACCCGGTTATTTTTTTTTTTTTTTTTTTTTTTTTT-3′)^40^ was pre-incubated with purified GajAB or GajAB + Gad1 in 20 µL DNA cleavage reactions containing 1 µM dsDNA, 1 µM GajAB or GajAB + Gad1, 1mM MgCl_2_, 20 mM Tris-HCl pH 9.0 for 1, 5, 10, 15, and 20 min at 37 °C. Following incubation, reactions were stopped with DNA loading buffer containing EDTA and 10 µL was analyzed on a 2% TB (Tris-borate) agarose gel. Gels were run at 250V for 40 min at 4 °C, then stained by rocking at room-temperature in TB buffer with 10 µg ml^-1^ ethidium bromide for 30 min. Gels were de-stained in TB buffer for 40 min and imaged with a ChemiDoc MP Imaging System.

### Construction of dCas9 and gRNA cassettes for integration in *B. subtilis thrC* site

The plasmid pJG_thrC_dCAS9_gRNA was constructed as previously described^33^. To insert new spacers, two fragments were amplified from pJG_thrC_dCAS9_gRNA and the new spacer was introduced into the overlap of primers designed for NEBuilder HiFi DNA Assembly (NEB, no. E2621). For the gRNA used to target *tad2*, the first fragment was amplified using primers JG528 and JG525, and the second using primers JG529 and JG524 (Supplementary table 15). The resulting assembled construct had the gRNA sequence “aagatgatgttcccaaacac”. For the gRNA used to target *had1*, the first fragment was amplified using primers JG389 and JG381, and the second using primers JG390 and JG382 (Supplementary table 15). The resulting assembled construct had the gRNA sequence “gcttgctaggattagtgtcc”. The gRNA sequence “ctatgattgatttttttagc” was used as a control. It was constructed as mentioned above, with primers JG389 and JG378, and JG390 and JG388 (Supplementary table 15). Shuttle vectors were propagated in *E. coli* DH5α with 100 μg ml^-1^ ampicillin selection. Plasmids were isolated from *E. coli* DH5α before transformation into the appropriate *B. subtilis* BEST7003 strains. The vectors containing the dCas9– gRNA sequences were cloned to *B. subtilis* strains containing the respective defense system, as well as to a control strain lacking the defense system.

### Knock-in of Had1 and Tad2 into phage SBSphiJ and Gad1 into phage SPbeta

The DNA sequence of *tad2*, together with the Phspank promoter, was amplified from the Tad2-containing pSG-*thrC*-Phspank plasmid using KAPA HiFi HotStart ReadyMix (Roche, cat #KK2601) with the primer pair tad2KIF and tad2KIr (Supplementary table 15). The DNA sequence of *had1*, together with its upstream intergenic region, was amplified from the genome of phage SBSphiJ4 with the primer pair had1KIF and had1KIR (Supplementary table 15). The backbone fragment with the upstream and downstream genomic arms (±1.2 kbp) for the integration site of *tad2* and *had1* was amplified from the plasmid used previously for knock-in of the *tad1* gene^33^, with the primer pair backboneKIF and backboneKIR (Supplementary table 15). The DNA sequence of *gad1*, together with its upstream intergenic region, was amplified from the genome of phage phi3T with the primer pair gad1KIF and gad1KIR (Supplementary table 15). The upstream and downstream genomic arms (±1.2 kbp) for the integration site of the *gad1* gene within the SPbeta genome were amplified from the genome of phage SPbeta using the primer pair gad1LFF and gad1LFR, and the primer pair gad1RFF and gad1LRR (Supplementary table 15). Cloning was conducted using the NEBuilder HiFi DNA Assembly cloning kit (NEB, no. E5520S) and transformed to DH5α-competent cells. The cloned vector was subsequently transformed into the *thrC* site of *B. subtilis* BEST7003.

The *tad2*, *had1* and *gad1* containing *B. subtilis* BEST7003 strains were then infected with phages SBSphiJ (*tad2* and *had1*) or SPbeta (*gad1*) with an MOI of 0.1 and cell lysates were collected. Tad2 lysate was used to infect a Thoeris-containing *B. subtilis* culture in two consecutive rounds with an MOI of 2 in each round (30 °C, 1mM IPTG). Had1 lysate was used to infect a Hachiman-containing *B. subtilis* culture in two consecutive rounds with an MOI of 2 in each round (25 °C). Gad1 lysate was used to infect a Gabija-containing *B. subtilis* culture in two consecutive rounds with an MOI of 2 in each round (25 °C). Several plaques were collected and screened using PCR for insertions. Phages with anti-defense genes were purified three times on *B. subtilis* BEST7003. Purified phages were verified again for the presence of *tad2*, *had1* and *gad1* using PCR amplifications.

### Identification of anti-defense homologs and phylogenetic reconstruction

Homologs of anti-defense genes were searched in the metagenomic gut virus (MGV)^45^ database using the “search” option of MMseqs release 12-113e3 with default parameters. Homologs of Gad1, Gad2 and Had1 were searched in the integrated microbial genomes (IMG) database^44^ using the blast option in the IMG web server. Gad1 and Gad2 homologs were searched using the default parameters, while Had1 homologs were searched using an e-value of 10 due to their short size. For Gad1 and Had1 this process was repeated iteratively for homologs that were found, until convergence. For Tad2, due to the multitude of homologs, homology search was done using the “search” option of MMseqs release 12-113e3 with default parameters, against ∼38000 prokaryotic genomes downloaded from the IMG database in October 2017.

For each family of anti-defense proteins, the unique (non-redundant) sequences were used for multiple sequence alignment with MAFFT version 7.402 ^65^ using default parameters. Phylogenetic trees were constructed using IQ-TREE version 1.6.5 ^66^ with the -m LG parameter. The online tool iTOL24 (v.5)^67^ was used for tree visualization. Phage family annotations were based on the prediction in the MGV database. The host phyla annotations were either based on the prediction in the MGV database, or the IMG taxonomy of the bacteria in which the prophage was found. Gabija and Thoeris defense systems were found in the bacterial genomes using DefenseFinder^39^ version 1.0.9 and database release 1.2.3.

### Preparation of filtered cell lysates

For generating filtered cell lysates, we used *B. subtilis* BEST7003 cells co-expressing Tad2 and the *B. cereus* MSX-D12 Thoeris system in which ThsA was inactivated (ThsB/ThsA_N112A_). Tad2 was integrated in the thrC locus as described above and expressed from an inducible Phspank promoter, and the Thoeris system was integrated in the amyE locus and expressed from its native promoter as described above. Controls included cells expressing only the ThsB/ThsA_N112A_ Thoeris system without Tad2, as well as cells lacking both the Thoeris system and Tad2. These cultures were grown overnight and then diluted 1:100 in 250 ml MMB medium supplemented with 1 mM IPTG and grown at 37°C, 200 rpm shaking for 120 min followed by incubation and shaking at 25 °C, 200 rpm until reaching an OD_600_ of 0.3. At this point, a sample of 40 ml was taken as the uninfected (time 0 min) sample, and SBSphiJ phage was added to the remaining 210 ml culture at an MOI of 10. Flasks were incubated at 25°C with shaking (200 rpm), for the duration of the experiment. 40 ml samples were collected at time points 75, 90, 105 min post-infection. Immediately upon sample removal (including time point 0 min), the 40 ml sample tubes were centrifuged at 4°C for 10 min to pellet the cells. The supernatant was discarded, and the pellet was flash frozen and stored at −80 °C.

To extract cell metabolites from frozen pellets, 600 µl of 100 mM Na phosphate buffer (pH 8.0) was added to each pellet. Samples were transferred to FastPrep Lysing Matrix B in a 2 ml tube (MP Biomedicals, no. 116911100) and lysed at 4 °C using a FastPrep bead beater for 2 × 40 s at 6 m s^−1^. Tubes were then centrifuged at 4 °C for 10 min at 15,000 *g*. Supernatant was then transferred to an Amicon Ultra-0.5 Centrifugal Filter Unit 3 kDa (Merck Millipore, no. UFC500396) and centrifuged for 45 min at 4 °C, 12,000 *g*. Filtered lysates were taken for *in vitro* ThsA-based NADase activity assay.

### Expression and purification of ThsA

*B. cereus* MSX-D12 *thsA* fused to a C-terminal TwinStrep tag was cloned into a pACYC-Duet1 plasmid (addgene plasmid #71147). The protein was expressed under the control of the T7 promoter together with a C-terminal Twin-Strep tag for subsequent purification. Expression was performed in 5 L LB medium supplemented with chloramphenicol (34 mg ml^−1^) in *E. coli* BL21(DE3) cells. Induction was performed with 0.2 mM IPTG at 15 °C overnight. The cultures were collected by centrifugation and lysed by a cooled cell disrupter (Constant Systems) in 100 ml lysis buffer composed of 20 mM HEPES pH 7.5, 0.3 M NaCl, 10% glycerol and 5 mM β-mercaptoethanol, 200 KU/100 ml lysozyme, 20 µg ml^−1^ DNase, 1 mM MgCl_2_, 1 mM phenylmethylsulfonyl fluoride (PMSF) and protease inhibitor cocktail (Millipore, 539134). Cell debris were sedimented by centrifugation, and the lysate supernatant was incubated with 2 ml washed StrepTactin XT beads (IBA, 2-5030–025) for 1 h at 4 °C. The sedimented beads were then packed into a column connected to an FPLC allowing the lysate to pass through the column at 1 ml min^−1^. The column was washed with 20 ml lysis buffer. ThsA was eluted from the column using elution buffer containing 50 mM biotin, 100 mM Tris pH 8, 150 mM NaCl and 1 mM EDTA. Peaks containing the ThsA protein were injected to a size-exclusion chromatography (SEC) column (Superdex 200_16/60, GE Healthcare, 28-9893-35) equilibrated with SEC buffer (20 mM HEPES pH 7.5, 200 mM NaCl and 2 mM DTT). Peaks were collected from the SEC column, aliquoted and frozen at −80 °C to be used for subsequent experiments.

### ThsA-based NADase activity assay

NADase reaction was performed in black 96-well half area plates (Corning, 3694). In each reaction microwell, purified ThsA protein was added to cell lysate, or to in vitro reactions of Tad2 with 1′′–3′ gcADPR, or to 100 mM sodium phosphate buffer pH 8.0. 5 µl of 5 mM nicotinamide 1,N6-ethenoadenine dinucleotide (εNAD^+^, Sigma, N2630) solution was added to each well immediately before the beginning of measurements, resulting in a final concentration of 100 nM ThsA protein in a 50 µl final volume reaction. Plates were incubated inside a Tecan Infinite M200 plate reader at 25 °C, and measurements were taken at 300 nm excitation wavelength and 410 nm emission wavelength. Reaction rate was calculated from the linear part of the initial reaction.

### Tad2 protein cloning, expression, and purification for biochemistry

The SPO1 *tad2* gene was cloned into the expression vector pET28-bdSumo as described previously^33^. Tad2 was expressed in *E. coli* BL21(DE3) by induction with 200 µM IPTG at 15 °C overnight. A 2L culture of Tad2 was harvested and lysed by a cooled cell disrupter (Constant Systems) in lysis buffer (50mM Tris pH 8, 0.25M NaCl, 10% Glycerol) containing 200KU 100 ml^-1^ lysozyme, 20ug ml^-1^ DNase, 1mM MgCl_2_, 1 mM phenylmethylsulfonyl fluoride (PMSF) and protease inhibitor cocktail. After clarification of the supernatant by centrifugation, the lysate was incubated with 5 ml washed Ni beads (Adar Biotech) for 1 h at 4 °C. After removing the supernatant, the beads were washed 4 times with 50 ml lysis buffer. Tad2 was eluted by incubation of the beads with 10 ml cleavage buffer (50 mM Tris pH 8, 0.25M NaCl, 10% Glycerol and 0.4 mg bdSumo protease) for 1 h at 23 °C. The supernatant, containing cleaved Tad2, was removed, and an additional 5 ml cleavage buffer was added to the beads and left overnight at 4 °C. The two elution solutions were combined, concentrated, and applied to a size exclusion (SEC) column (HiLoad_16/60_Superdex75 prep-grade, Cytiva) equilibrated with 50 mM Tris pH 8, 100 mM NaCl. Pure Tad2 migrating as a single peak was pooled and flash frozen in aliquots using liquid nitrogen and stored at −80 °C.

### Incubation of purified 1′′–3′ gcADPR with Tad2

2.4 µM purified Tad2 was incubated with 600 nM of 1′′–3′ gcADPR in 100 mM Na Phosphate buffer, pH 8.0 for 10 min, at 25°C, followed by an additional 10 min incubation at either 95°C or 25°C. Following incubation, samples were left on ice for 1 min. The samples were then used for ThsA-based NADase activity assays as described above.

### Analytical SEC analysis of apo and ligand bound Tad2

50 µl of Tad2 (158 µM) was incubated with 30 µl of 1 mM 1’’–3’ gcADPR at 25 °C, in 0.1 M NaCl, 50 mM TrisHCl pH 8.0, for 20 min. The incubated mixture, and an apo protein incubated with an identical buffer without the ligand, were then loaded on a size exclusion Superdex_200_10/300 analytical column (PBS buffer) and monitored for absorption at both 260_nm_ and 280_nm_. The oligomeric nature of Tad2 apo protein was evaluated by comparing the retention time of apo-Tad2 to that of five internal standard proteins (Ribonuclease 13.7 kD, Chymotrypsin 25 kD, BSA 66 kD, Aldolase 160 kD, and apoferritin 443 kD) using a size exclusion Superdex_200_10/300.

### Surface Plasmon Resonance (SPR) measurements of Tad2 binding to 1′′–3′ gcADPR and cADPR

Association and dissociation of 1′′–3′ gcADPR or cADPR to/from Tad2 were monitored by surface plasmon resonance with a BIAcore S200 apparatus (Cytiva, Sweden). Tad2 was immobilized on a CM5 S Series chip (Cytiva, Sweden) by amine coupling chemistry using the following protocol: chip activation was made with a freshly prepared mixture of N-hydroxysuccinimide (50 mM in water) and 1-ethyl-3-(3-dimethylaminopropyl) carbodiimide (195 mM in water) for 7.5 min in DPBS (Sartorius, SKU 02-023-5A) (flow rate of 10 μL/min). DPBS served as the running buffer. Tad2 protein (5 μg ml^−1^ in 300 mM sodium acetate buffer, pH 3.8) was injected for 5 min (flow rate 10 μL/min) and the remaining activated carboxylic groups were blocked by the injection of 1 M ethanolamine hydrochloride, pH 8.6, for 5 min (flow rate 10 μL/min). A total of 1,800 RU of Tad2 were immobilized by this method. Associations of 1′′–3′ gcADPR and cADPR with Tad2 were monitored by injecting 1′′–3′ gcADPR/cADPR at multiple concentrations for 2 min at 25 °C and flow rate of 50 μL/min. No regeneration step was required after ligand binding. Sensograms were fitted to steady state model (S200 evaluation software 1.1).

### Protein cloning, expression, and purification for gcADPR production and protein crystallization

Synthetic DNA fragments (Integrated DNA Technologies) of cmTad1, AbTIR^TIR^ (Δ1-156), cbTad1, and SPO1 Tad2 genes were cloned into a custom pET expression vector containing an N-terminal 6×His-SUMO tag and an ampicillin resistance gene by Gibson assembly, as previously described^68^. Three colonies of BL21(DE3) RIL *E. coli* transformed with these plasmids onto MDG agar plates (1.5% agar, 2 mM MgSO_4_, 0.5% glucose, 25 mM Na_2_HPO_4_, 25 mM KH_2_PO_4_, 50 mM NH_4_Cl, 5 mM Na_2_SO_4_, 0.25% aspartic acid, 2–50 μM trace metals) were picked into 30 ml MDG starter culture and shaken overnight at 230 rpm and 37 °C. 1L M9ZB expression medium (2 mM MgSO_4_, 0.5% glycerol, 47.8 mM Na_2_HPO_4_, 22 mM KH_2_PO_4_, 18.7 mM NH_4_Cl, 85.6 mM NaCl, 1% Cas-amino acids, 2–50 μM trace metals, 100 μg ml^−1^ ampicillin and 34 μg ml^−1^ chloramphenicol) was seeded with 15 ml starter culture and grown at 230 rpm and 37 °C to an OD_600_ of 2 before induction of expression with 0.5 mM IPTG and incubation at 230 rpm and 16°C for 16 hours. For AbTIR^TIR^ expression, nicotinamide-supplemented 2YT expression medium (1.6% w/v tryptone, 1% w/v yeast extract, 342 mM NaCl, 10 mM NAM, 100 μg ml−1 ampicillin, 34 μg ml−1 chloramphenicol) was used instead. Cells were harvested by centrifugation, resuspended in lysis buffer (20 mM HEPES-KOH pH 7.5, 400 mM NaCl, 30 mM imidazole, 10% glycerol, 1 mM DTT), lysed by sonication, and clarified by centrifugation at 25,000*g* for 20 min. Lysate was passed over 8 ml Ni-NTA resin (Qiagen), washed with 70 ml wash buffer (20 mM HEPES-KOH pH 7.5, 1 M NaCl, 30 mM imidazole, 10% glycerol, 1 mM DTT), eluted with lysis buffer supplemented to 300 mM imidazole, and dialyzed in 14 kDa dialysis tubing in dialysis buffer (20 mM HEPES-KOH pH 7.5, 250 mM KCl, 1 mM DTT) overnight at 4°C with purified human-SENP2 for 6× His-SUMO tag cleavage. For crystallography, proteins were further purified on a Superdex 75 16/600 size-exclusion chromatography column (Cytiva). Final samples were concentrated to >20 mg mL^-1^, flash frozen, and stored at −80°C.

### 1′′–2′ gcADPR and 1′′–3′ gcADPR production and purification

gcADPR molecules were produced as described previously^33^. For 1′′–2′ gcADPR production, purified AbTIR^TIR^, a bacterial enzyme that efficiently converts NAD^+^ to 1′′–2′ gcADPR^69^, was used to set up 300μl reactions (50 mM HEPES-KOH, 150 mM NaCl, 20 mM NAD^+^, 40 μM AbTIR^TIR^). For 1′′–3′ gcADPR production, Purified ThsB’^33^ was used to set up 50 ml reactions (50 mM HEPES-KOH pH 7.5, 150 mM NaCl, 2 mM NAD^+^, 16 μM ThsB’). Reactions were carried out at RT for 24-48 hours before boiling at 95°C for 10 min, pelleting at 13,500 *g* for 20 min, and filtering through a 10 kDa MWCO filter (Amicon). For the AbTIR^TIR^ reaction, filtrate was diluted to 30 ml with PBS followed by addition of 200 μM purified cmTad1. For the ThsB’ reaction, 10 μM purified cmTad1 was added directly to 50 ml of filtrate. Mixtures were then incubated for 1 hour at RT to allow cmTad1–gcADPR complex formation. The complexes were washed by successive concentration and dilution in a 10 kDa MWCO filter first with five 1:20 dilutions in PBS followed by five 1:20 washes in water. Complexes were concentrated to >3 mM before final release and extraction of gcADPR by boiling at 95 °C for 10 min, pelleting at 13,500 *g* for 20 min, filtering through a 3 kDa MWCO filter, and collecting the filtrate. Final purity and concentration were assessed by HPLC.

ThsB′ was cloned and purified similarly as described above, except that it was cloned into a custom pET vector containing a C-terminal 6× His tag and a chloramphenicol resistance gene, transformed into BL21(DE3) cells, grown in the presence of chloramphenicol only, expressed in 2YT medium with 10 mM NAM, and concentrated to 4 mg ml^−1^ after dialysis before flash-freezing and storage.

### Protein crystallization and structural analysis

cbTad1 and SPO1 Tad2 crystals were grown by the hanging-drop method in EasyXtal 15-well trays (NeXtal) at 16°C. Hanging-drops were set using 1 μl of diluted protein solution (5-10 mg mL^-1^ protein, 20 mM HEPES-KOH pH 7.5, 80 mM KCl, 1 mM TCEP) and 1 μl reservoir solution over a 400μl well of reservoir solution. Proteins were crystallized and cryoprotected under the following conditions before being harvested by flash-freezing in liquid nitrogen: (1) cbTad1 complexed with 1′′–3′ gcADPR: Crystals were grown for 3 weeks in drops supplemented with 500 μM ThsB′-derived 1′′–3′ gcADPR using reservoir solution containing 0.2 M MgCl_2_, 0.1 M Tris-HCl pH 8.5, and 3.4 M 1,6-hexanediol. (2) SPO1 Tad2 in the apo state: Crystals were grown for 1 week using reservoir solution containing 0.1 M 2-(*N*-morpholino) ethanesulfonic acid pH 6.5, 10% (v/v) 1,4-dioxane, and 1.6 M ammonium sulfate before being cryoprotected with reservoir solution supplemented with 25% (v/v) glycerol. (3) SPO1 Tad2 complexed with 1′′–3′ gcADPR: Crystals were grown for 1 week in drops supplemented with 500 μM ThsB′-derived 1′′–3′ gcADPR using reservoir solution containing 0.2 M magnesium nitrate and 18% (w/v) PEG 3350 before being cryoprotected with reservoir solution supplemented with 20% (v/v) ethylene glycol and 1 mM 1′′–3′ gcADPR. (4) SPO1 Tad2 complexed with 1′′–2′ gcADPR: Crystals were grown for 1 week in drops supplemented with 500 μM AbTIR^TIR^-derived 1′′–2′ gcADPR using reservoir solution containing 0.1 M Tris-HCl pH 8.5, 12% (v/v) glycerol, and 1.5 M ammonium sulfate before being cryoprotected with reservoir solution supplemented with 15% (v/v) ethylene glycol and 1 mM 1′′–2′ gcADPR. X-ray diffraction data were collected at the Advanced Photon Source (beamline 24-ID-C), and data were processed using the SSRL autoxds script (A. Gonzalez, Stanford SSRL). Phases were determined by molecular replacement in Phenix using either previously determined cbTad1 structures^33^ (PDB 7UAV, 7UAW) or sequence-predicted SPO1 Tad2 truncated structures from ColabFold v1.5.2 ^70^. Model building was performed in WinCoot, refinement was performed in Phenix^71, 72^, statistics were analyzed as presented in Extended Data Table 1^73–75^, and structure figures were produced in PyMOL. Final structures were refined to stereochemistry statistics for Ramachandran plot (favoured/allowed), rotamer outliers and MolProbity score as follows: cbTad1–1′′–3′-gcADPR, 95.88/4.12%, 0.92%, 1.47; SPO1 Tad2 apo, 96.15/2.56%, 0.74%, 1.59; SPO1 Tad2–1′′–3′-gcADPR, 99.18/.82%, 1.5%, 1.33; SPO1 Tad2–1′′–2′-gcADPR, 99.36/0.64%, 2.19%, 1.27.

### HPLC analysis of Tad2 incubation with 1′′–3′ gcADPR

Reactions to analyze cleavage of 1′′–3′ gcADPR were performed in 120 μl reactions consisting of 50 mM Tris-HCl pH 7.5, 100 mM KCl, 5 mM MgCl_2_, 1 mM MnCl_2_, 50 μM gcADPR isomer, and either buffer or 1 μM Tad2. As a control, 1′′–3′ gcADPR was also incubated with 1 μl Cap-Clip Acid Pyrophosphatase (known to cleave diphosphate linkages on mRNA caps, Fisher Scientific), using the manufacturer’s recommended reaction conditions. Reactions were incubated at 37 °C for 1 hour before filtration using a 3 kDa MWCO filter. Filtered reactions were analyzed using a C18 column (Agilent Zorbax Bonus-RP 4.6 x 150 mm) heated to 40 °C and run at 1 ml/min using a buffer of 50 mM NaH_2_PO_4_-NaOH pH 6.8 supplemented with 3% acetonitrile.

## Acknowledgements

We thank the Sorek laboratory members for comments on the manuscript and fruitful discussion. We also thank Y. Peleg and S. Albeck from the Center for Structural Proteomics within the Weizmann Institute of Science for assistance with protein expression, Y. Fridmann-Sirkis from the Life Sciences Core Facilities of the Weizmann Institute for help with SPR analysis, and H. Keren-Shaul and D. Pilzer from the Life Sciences Core Facilities of the Weizmann Institute for help with PacBio sequencing. R.S. was supported, in part, by the European Research Council (grant no. ERC-AdG GA 101018520), Israel Science Foundation MAPATS Grant 2720/22), the Deutsche Forschungsgemeinschaft (SPP 2330, Grant 464312965), the Ernest and Bonnie Beutler Research Program of Excellence in Genomic Medicine, and the Knell Family Center for Microbiology. E.Y. is supported, in part, by the Israeli Council for Higher Education (CHE) via the Weizmann Data Science Research Center. P.J.K. was supported, in part, by the Pew Biomedical Scholars programme and The Mathers Foundation. S.J.H. is supported through a Cancer Research Institute Irvington Postdoctoral Fellowship (no. CRI3996). X-ray data were collected at Northeastern Collaborative Access Team beamlines 24-ID-C and 24-ID-E (no. P30 GM124165), including use of a Pilatus detector (no. S10RR029205), an Eiger detector (no. S10OD021527) and the Argonne National Laboratory Advanced Photon Source (no. DE-AC02-06CH11357).

## Author Contribution

The study was conceptualized and designed by E.Y., A. Leavitt and R.S. E.Y. built and executed the computational pipeline and analyzed the data. A. Leavitt isolated the phages and conducted all the *in vivo* experiments unless stated otherwise. A. Lu and P.J.K. preformed the structural analysis of Tad1 and Tad2. C.A. and G.A. preformed the biochemical experiments with cell lysates and led the mechanistic characterization of the Tad2 activity. I.O. designed and conducted all the phage knock in experiments and the knockout of *gad2* from phage SPbetaL7. J.G. designed and generated the knock down clones. DNA cleavage experiments were performed by S.P.A., S.E.M., and P.J.K. S.J.H. helped with the structural analysis and with the characterization of the Tad2 activity. The study was supervised by G.A. and R.S. The manuscript was written by E.Y. and R.S. All authors contributed to editing the manuscript and support the conclusions.

